# Computing Hubs in the Hippocampus and Cortex

**DOI:** 10.1101/513424

**Authors:** Wesley Clawson, Ana F. Vicente, Maëva Ferraris, Christophe Bernard, Demian Battaglia, Pascale P Quilichini

**Affiliations:** Aix Marseille Univ, Inserm, INS, Institut de Neurosciences des Systèmes, Marseille, France

## Abstract

Neural computation, which relies on the active storage and sharing of information, occurs within large neuron networks in the highly dynamic context of varying brain states. Whether such functions are performed by specific subsets of neurons and whether they occur in specific dynamical regimes remains poorly understood. Using high density recordings in the hippocampus, medial entorhinal and medial prefrontal cortex of the rat, we identify computing substates, or discrete epochs, in which specific computing hub neurons perform well defined storage and sharing operations in a brain state-dependent manner. We retrieve a multiplicity of distinct computing substates within each global brain state, such as REM and nonREM sleep. Half of recorded neurons act as computing hubs in at least one substate, suggesting that functional roles are not firmly hardwired but dynamically reassigned at the second timescale. We identify sequences of substates whose temporal organization is dynamic and stands between order and disorder. We propose that global brain states constrain the language of neuronal computations by regulating the syntactic complexity of these substate sequences.

---

Information processing in the brain can be approached on three different levels: biophysical, algorithmic and behavioral (*1*). The algorithmic level, which remains the least understood, describes the way in which emergent functional computations can be decomposed into simpler processing steps, with architectures mixing serial and massively parallel aspects (*2*). At the lowest level of individual system components - here, in single neurons, such building blocks of distributed information processing can be modeled as primitive operations of storing, transferring, or non-linearly integrating information streams (*3*).

In resting state conditions, both BOLD and EEG signals are characterized by discrete epochs of functional connectivity or topographical stability, defined as resting state networks and microstates, respectively (*4*-*5*). The transitions between these large-scale epochs are neither periodic nor random but occur through a not yet understood syntax, which is fractal and complex (*5*). Does such organization at the macroscopic scale (whole brain and networks of networks for resting state networks and microstates, respectively) also exist at the microscopic scale? Said differently, is neuronal activity at the microcircuit level organized in discrete epochs associated to different “styles” of information processing? Our first goal is to determine whether information processing at the local neuronal circuit level is structured into discrete sequences of substates, and whether such sequences have an observable syntax, whose complexity could be a hallmark of computation. Here we focus on low-level computing operations, performed by individual neurons such as basic information storage and sharing (*6*-*7*). To reduce external perturbations, such as sensory inputs, and to establish if primitive processing operations and their temporal sequences are brain state-dependent, we study two conditions: anesthesia and natural sleep, which are characterized by alternating stable brains states, theta (THE)/slow oscillations (SO) and rapid eye movement (REM)/nonREM sleep, respectively. We consider the CA1 region of the hippocampus, the medial entorhinal cortex (mEC) and the medial prefrontal cortex (mPFC) to determine whether algorithmic properties are shared between regions with different cytoarchitectures.

The second goal is to determine whether primitive processing operations are localized, or on the contrary, distributed within the microcircuit as proposed for attractor neural networks (8) and liquid state machines (*9*). This raises two key questions: Are certain operations driven by a few key neurons, similar to hub cells in a rich club architecture (*10*)? and Do neurons have pre-determined computing roles, such as ‘sharer’ or ‘storer’ of information’, as well as rigidly prescribed partners in their functional interactions? Said differently - is information routed through a hardwired ‘neuronal switchboard system’ like in early days of telephony? Or dynamically via different addressable nodes like in decentralized peer-to-peer services?

Here we demonstrate the existence of a multiplicity of distinct computing substates at the microcircuit level within each of the probed global brain states in both anesthesia and natural sleep. The low-level algorithmic roles played by individual neurons change from one substate to the other and appear largely independent from the underlying cytoarchitecture, with roughly half of the recorded neurons acting as transient computing hubs. Furthermore, we reveal complexity not only at the level of information processing within each substate but also at the level of how substates are organized into temporal sequences, which are neither regularly predictable nor fully random. Substate sequences display an elaborate syntax in all the probed anatomical regions, whose complexity is systematically modulated by changes in global brain states.

Taken together, our findings suggest a more distributed and less hierarchical style of information processing in neuronal microcircuits, more akin to emergent liquid state computation than to pre-programmed processing pipelines.

## RESULTS

### Analysis design

Neurons were recorded simultaneously from the CA1 region of the hippocampus and the medial entorhinal cortex (mEC) under anesthesia (18 recordings from 16 rats), and from the CA1 region and the medial prefrontal cortex (mPFC) during natural sleep (6 recordings from 3 rats, see Figures 1A, S1 and S2 for more details on recordings). We focus on two elementary processing functions: *information storage*, i.e. how much information a neuron buffers over time that it has previously conveyed, as measured by the active information storage (*3*); and *information sharing*, i.e. how much a neuron’s activity information content is made available to other neurons, as measured by mutual information (see e.g. in *7*). We use the term *feature* to discuss the metrics we use; i.e. firing, information storage or sharing (Figure 1B-C). We use the same analysis design for all features. The `FeatureVector(`*t_a_*) contains the values for the descriptive features as measured in window *t_a_* (Figure 1B). For example, for firing features, if 20 cells are recorded, `FeatureVector(`*t_a_*) contains 20 values, representing the firing density of each neuron during window *t_a_*. We first correlate feature vectors for a given window pair (*t_a_*, *t_b_*). Here, a high correlation value means that the two feature vectors are very similar to one another, i.e. that the features measured at *t_a_* are also found at *t_b_*. After, we build a *feature similarity matrix*, a collection of correlation values between feature vectors for all window pairs, organized in time (Figure 1D). A block along the diagonal indicates a stable state for a given feature, e.g., a period over which units fire, store or share information in a consistently preserved pattern. The axes of the similarity matrix represent time, and repetitions of a block structure along a horizontal or vertical line mean that a stable state for a given feature is reoccurring over time. We then use a simple clustering technique to extract different stable states, which we call *substates*, and display their switching behavior during the recording session (Figure 1D). Finally, we define *computing hubs* as neurons that more heavily participate to the buffering (storage hubs) or the funneling (sharing hubs) of information streams (Figure 1D, see Material and Methods). This notion of computing hub generalizes previously introduced notions of “hubness” (25, 26) beyond the ability to synchronize firing toward more general types of influence on information processing.

**Figure 1.**
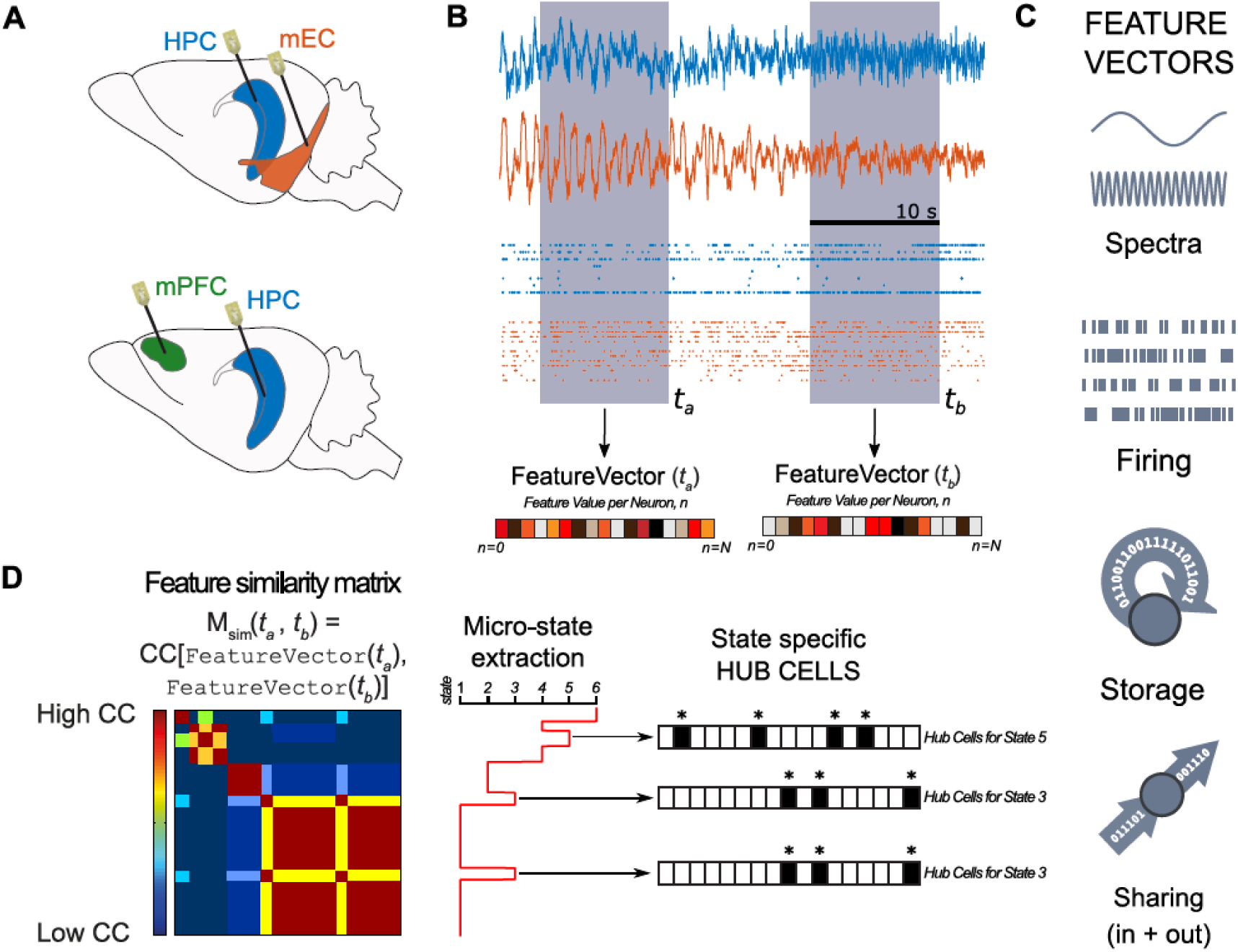
Unsupervised extraction of states and hubs. **A**. Cartoon representing the approximate recording locations (mEC and CA1, mPFC and CA1) during 2 experiment types in anesthesia and sleep. **B**. Example LFP trace taken from the 32 channels in CA1 (blue) and 32 channels in mEC (orange). Below are examples of isolated unit activity taken from the same recording. For each time window (*t*), we extract different features represented by the `FeatureVector`*(t)*, which has a feature value for each channel or single unit recorded. **C.** We consider four features: spectral band averaged powers (from LFP channels); single unit firing rates; information storage and information sharing. **D.** Left panel: A cartoon representation of *M_sim_*. To extract substates and their temporal dynamics, we construct a *feature similarity* matrix *M_sim_* in which the entry *M_sim_(t_a_, t_b_)* measures Pearson correlation between the vectors `FeatureVector`*(t_a_)* and `FeatureVector`*(t_b_)*. Time flows from the top-left corner horizontally to the top-right corner and vertically to the bottom-left corner. A block (square) along the diagonal in the resulting image identifies a period of feature stability, i.e. a substate. A block appearing several times horizontally or vertically indicates that a feature is repeated several times. Middle panel: Unsupervised clustering identifies the different substates (indicated by a number) and their temporal dynamics (the vertical axis corresponds to that of the similarity matrix). Right panel: We identify computing hub cells, i.e. neurons that display exceptionally high values for a given feature, associated with given substates. Note that reoccurring states have the same hub cells (state 3 in this example).

### Identification of brain global states

Unsupervised cluster analysis of the spectral features of the fields recorded in the various brain regions allowed a clear identification of typical global oscillatory patterns (Figure S3), which we call *global brain states*. In the following, all brain states are identified by the clustering analysis of field recordings performed in the CA1 region (stratum oriens to stratum lacunosum moleculare). Unsupervised clustering identified two states for anesthesia corresponding to epochs dominated by slow (SO state) and theta (THE state) oscillations; and two states during sleep corresponding to REM vs nonREM episodes.

### Brain state-dependent firing substates

As subsets of cells tend to fire spontaneously together in stereotypical patterns (*11*-*12*), we first analyzed neuronal firing assemblies. Figure S4 shows that the firing rate, the burst index and entrainment by the phase of the ongoing oscillations were brain region- and brain state-dependent as previously reported (*13*-*14*). A simple visual inspection of firing behavior revealed the probable existence of different firing sets, as some neurons tended to fire together during certain epochs; with these epochs repeating themselves over time (Figure S5). To quantify this observation, we constructed the feature vectors `Firing`(*t_a_*), whose entries are given by the average firing rate of each neuron within the window of analysis *t_a_*. The complex block structure of the similarity matrix revealed a repertoire of state transitions much richer than the one associated to global brain states (Figure 2). In this example, unsupervised clustering revealed a total of six firing substates in mEC (Figure 2A) and five in mPFC (Figure 2D) during THE and REM, respectively, for the two animals. Figure 2B demonstrates that a given brain state was characterized by the switching between different firing substates. Figure 2E shows that a subset of firing substates was shared between brain states, and importantly that the switch from one firing substate state to another did not necessarily coincide with a change in the brain global state (and vice versa). Quantification over all recordings revealed that firing substates occurred 87% of the time during either one of the possible global brain states (Figures 2C and 2F). Substates were found in the mEC, CA1 and mPFC and we found an average of ∼5 substates for all brain regions and brain states (Table 1). These results reveal that, although field recordings show stereotyped oscillatory behavior during a given brain state, the firing behavior of neurons display a richer dynamic repertoire. Their activity is compartmentalized in a small number of firing substates, with discrete switching events from one substate to another. The firing substates are brain state and brain region specific, and they are not strictly entrained by the global oscillatory state.

**Figure 2.**
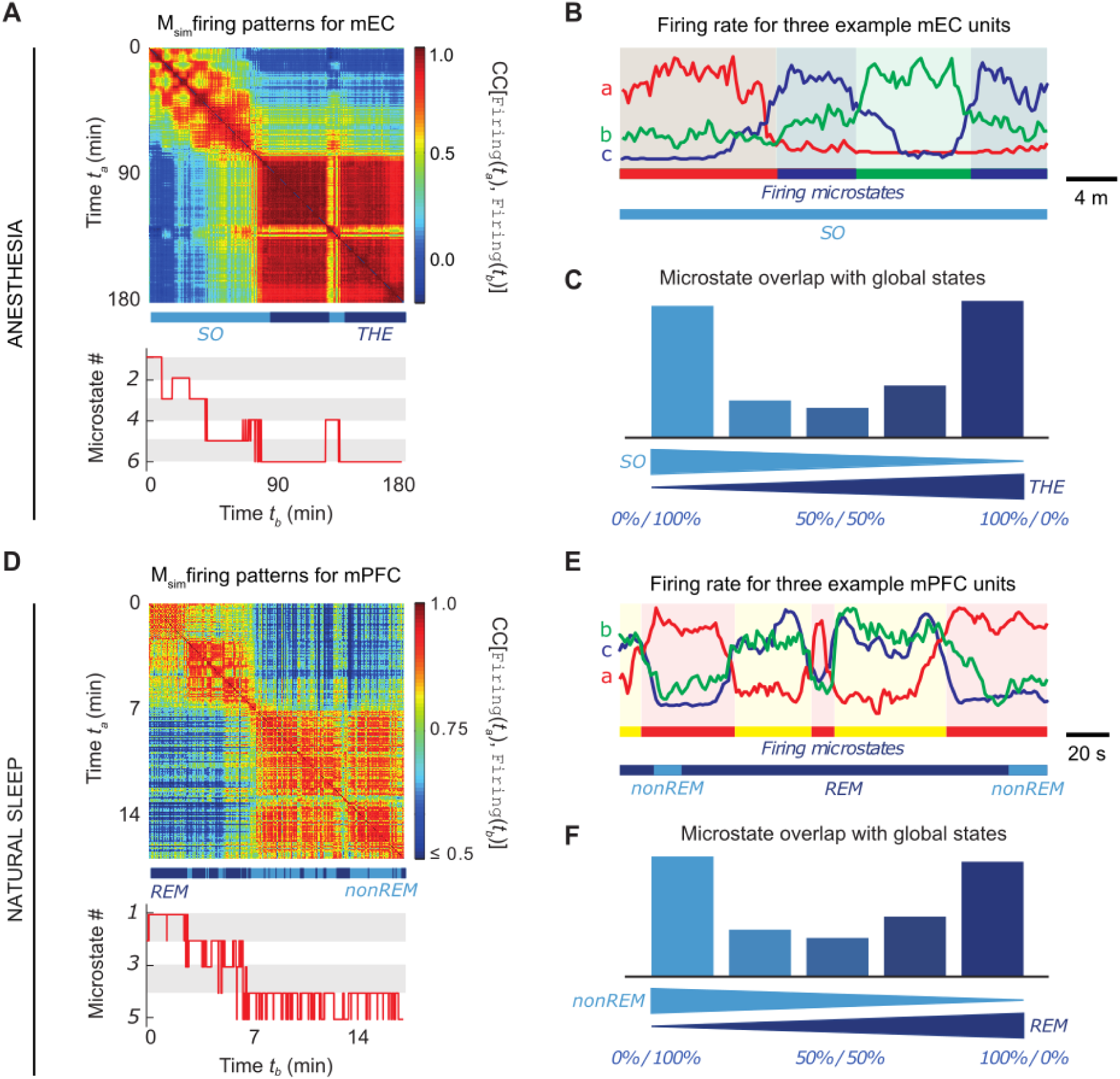
Firing substates. Examples of similarity matrices *M_sim_* obtained from `Firing`*(t)* at different times in mEC during anesthesia **(A)** and in mPFC during natural sleep **(D)**, measured in two animals. The bar below *M_sim_* indicates the transitions occurring between THE/REM (dark blue) and SO/nonREM (light blue). Although there were only two global brain states, six **(A)** and five **(D)** firing substates were identified. Panels **(B)** and **(E)** show examples of the firing density of three neurons (a, b and c) recorded in mEC and mPFC, respectively, with amplitude normalized for visualization. Neurons tended to fire in specific substates, indicated here with a color code. These examples also illustrate the switching between different firing substates inside a given global oscillatory state, and their overlap across different global oscillatory states. The analysis of all recordings revealed that a majority of firing substates tended to occur during a preferred global oscillatory state, as indicated by the bimodal histograms during anesthesia **(C)** and natural sleep **(F)**, respectively.

**Table 1.** Number of states and their oscillatory mode specificity

| State type | Oscillatory mode specificity | Median # states |
| --- | --- | --- |
| *Global spectral states* | *1.0* | *2* |
| Firing substates | 0.88 (0.78, 0.95) | 5 |
| Storage substates | 0.80 (0.69, 0.88) | 4 |
| Sharing substates | 0.86 (0.78, 0.95) | 4 |

*Median (1<sup>st</sup>, 3<sup>rd</sup> quartile), all regions and conditions*

### Storage of information is dynamic within a brain state

At any given time, neuronal activity conveys an amount of information that can be measured by Shannon entropy. We first focused on active information storage, which measures the fraction of information carried by a neuron *i* at a time *t* that was present in the past activity history of *i* itself (Figure S6A). For storage features, we extract several substates (6 for the mEC in the animal shown in Figure 3A, and 7 for CA1 in the animal shown in Figure 3D), with an average of ∼4 states across all animals (Table 1). As before, there was no strict alignment between brain state transitions and storage substate transitions (Figures 3A, B and C, E). Yet, brain state specificity of storage states was 80% for all regions (Figures 3C and F and Table 1).

**Figure 3.**
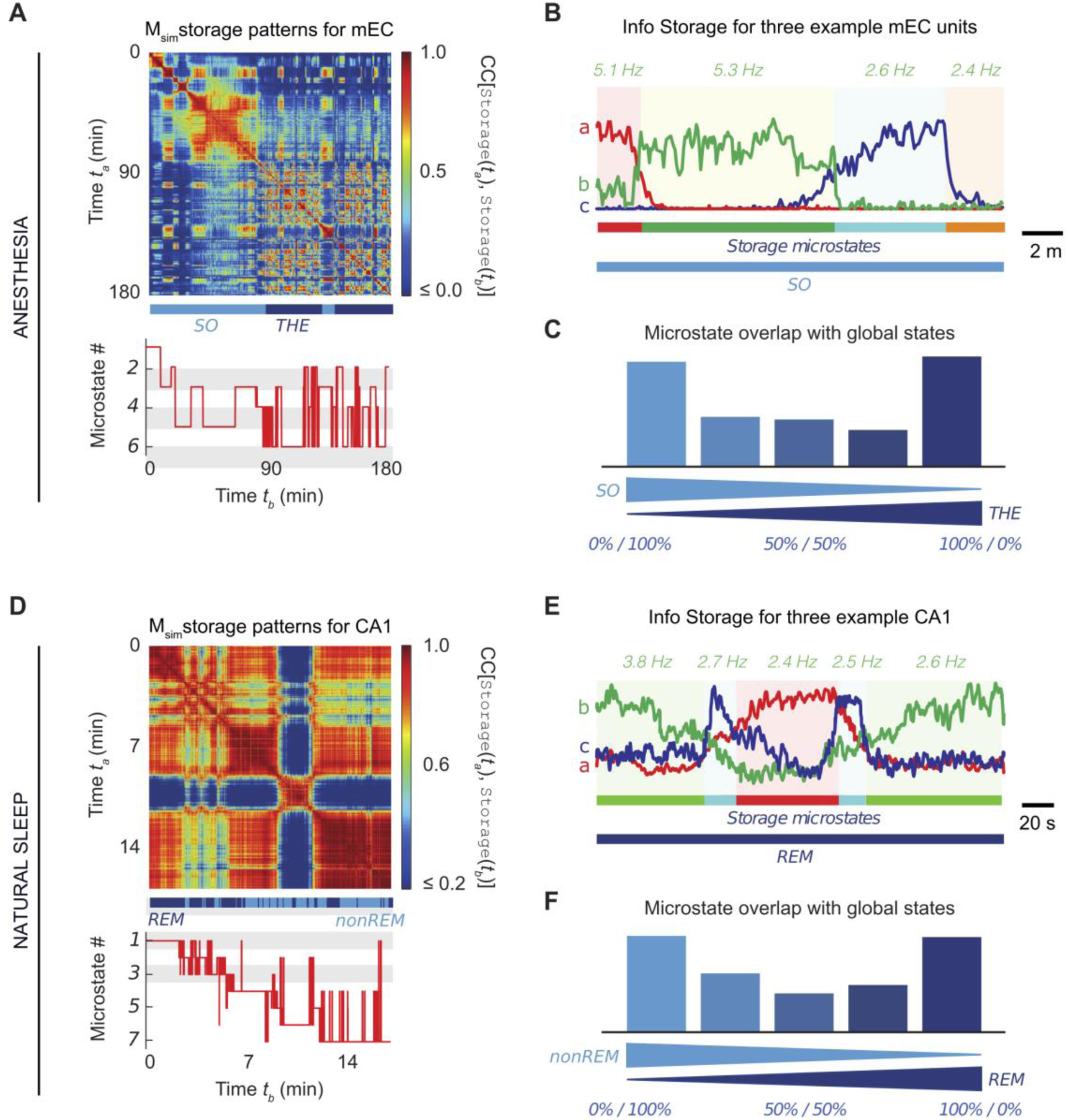
Information storage substates. Examples of similarity matrices *M_sim_* obtained from `Storage`*(t)* at different times in mEC during anesthesia **(A)** and CA1 during natural sleep **(D)**. As for firing substates, we identified more storage substates (6 and 7, respectively, in the shown examples) than global oscillatory states. We show in panels **(B)** and **(E)** that the participation of three individual neurons to information storage (indicated in arbitrary units for visualization) was substate-dependent. The values reported above the plots correspond to the average firing rate of the neuron *b* (green color) during the corresponding epochs within consistent storage substates. The analysis of all recordings showed that storage substates tended to occur during a preferred global oscillatory substate, as indicated by the bimodal histograms for anesthesia **(C)** and for natural sleep **(F)**.

Under anesthesia, the absolute storage values were stronger in mEC than in CA1, particularly in layers 3 and 5 of mEC (Figure S7). During natural sleep, however, storage values for CA1 were two orders of magnitude larger than during anesthesia and were as strong as in mPFC (Figure S7). Storage tended to be weaker for all probed regions and layers in THE with respect to SO during anesthesia, but not during natural sleep (Figure S7). Therefore, information storage is dynamically distributed in discrete substates and is brain state- and region-dependent. In particular, the involvement in storage of a neuron could vary substantially along time without being necessarily paralleled by a comparable change in firing rate (Figures 3B and 3E).

### Information sharing is dynamic within a brain state

A primitive processing operation complementary to information storage is information sharing, providing a pseudo-directed metric of functional connectivity between any two circuit units (*7*). For each neuron *i* we quantified both “shared-in” (*i* acts as a sharing target, with information shared from *j* neurons’ past activity, Figure S6B) and “shared-out” information (*i* acts as a sharing source and information is shared to *j* neurons’ future activity). We first constructed the feature vector `Sharing`(*t_a_*) containing the total amount of information funneled through each given neuron (integrated in- and out-sharing strengths, represented by big arrows in Figure 4A), irrespective of whom the information was being shared with. Since in- and out-sharing strengths were strongly correlated (average Pearson correlation >0.9), we ignored the distinction between in- and out-sharing and speak generically of sharing substates. Representative sharing similarity matrices and state sequences are shown in Figure 4B (top) for mEC during anesthesia and mPFC during sleep and in Figure S8 for CA1. Here, we studied only information sharing within regions, because the number of pairs of simultaneous units in different regions that showed significant sharing was too small to reach robust conclusions. We found ∼4 sharing substates on average across animals. Sharing states displayed an 86% specificity for a given brain state (Figure 4D, Table 1).

**Figure 4.**
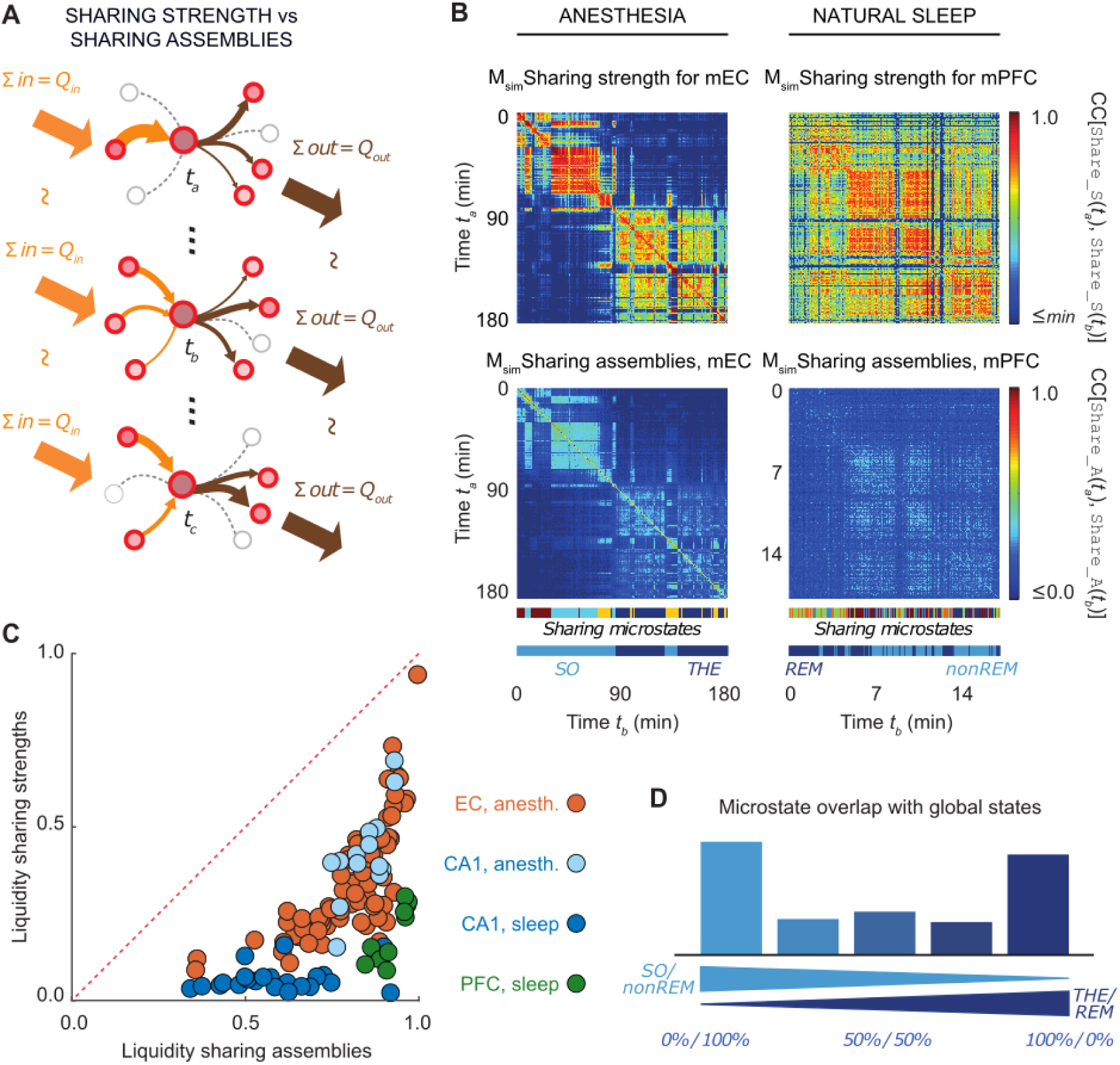
Information sharing substates. The cartoon in panel **A** shows an example of sharing assembly for a given sharing hub neuron across 3, non-sequential occurrences of the same substate. The total strength of in- and out-going sharing is equal (large, external arrows) during *t_a_, t_b_,* and *t_c_* while the assembly changes (smaller, internal arrows). The changing size of internal arrows represent the sharing strength of that particular functional connection between the sharing hub and its source and target neurons. **(B)** Similarity matrices *M_sim_* for sharing strengths `Sharing_S`*(t)* (top) and sharing assemblies `Sharing_A`*(t)* (bottom), in mEC during anesthesia (left) and mPFC during natural sleep (right). We identified a multiplicity of substates within each global oscillatory state as shown by the colored bars below the feature similarity matrices. The similarity matrices for sharing strengths and assemblies have a matching block structure. However, sharing strengths were very stable within a substate (red-hued blocks), while sharing assemblies were highly volatile (light blue-hued blocks). This is quantified for each sharing assembly substate by a *liquidity* coefficient (**C**). As shown in **(C)**, for all observed sharing substates across all regions and global oscillatory states in all animals, the liquidity of sharing assemblies was much larger than the one of sharing strengths. Finally, **(D)** demonstrates that most sharing substates occurred preferentially during a preferred global oscillatory state for both anesthesia and natural sleep combined (see Figure S7 for separated histograms for the two conditions).

During anesthesia, we measured a stronger absolute sharing values in CA1 than in mEC, a pattern reversed with respect to storage values, particularly in stratum radiatum (SR) and stratum pyramidale (SP) of CA1, even though mEC layer 5 had a sharing strength comparable to CA1s SR and SP (Figure S9). During natural sleep, the participation to information sharing of SO in CA1 increased by an order of magnitude and was as large as the one of mPFC, notably layer 4 (Figure S9). As for storage, the involvement of a neuron in sharing could vary along time even without corresponding variations of its firing rate (Figures S8B and S8E).

### Sharing assemblies are “liquid”

The previous analysis is focused on sharing strengths at the single cell level. We then determined with which neurons sharing cells were exchanging information, i.e. the detailed network neighborhood of sharing, or *sharing assembly* (cartoon networks in Figure 4A). Two striking features were apparent. First, both the block structure of the sharing assemblies and the state transition sequences are nearly matching the sharing strength ones (Figure 4B), as evidenced by a relative mutual information value of 98% on average. Second, in contrast to sharing strengths, the blocks in the sharing assembly similarity matrix were of a light blue color, indicating a strong variability of sharing assemblies within a given substate. This phenomenon was quantified by *liquidity* analysis, with liquidity being a measure bounded between 0 and 1, where a value of 0 represents an absence of internal variability within a substate and a value of 1 representing completely random variability (see *Materials and Methods*). The liquidity values of sharing assemblies for all sharing substates throughout all recordings lied below the diagonal (Figure 4C). This result can be better understood considering the toy examples of Figure 4A. The cartoons represent snapshots at three different, non-sequential times of a given hub neuron in its sharing network environment. The three considered time frames all fall within the same substate, therefore the overall in- and out-sharing strengths, represented by the orange and grey arrows respectively, are constant (meaning stability). However, the sources and targets of the funneled information can widely vary in time (meaning instability). Although the sum of in- going and out-going information remained overall constant within each sharing substate, information was shared over different cell assemblies from one time period to the next. All three brain regions displayed remarkable liquidity in sharing assemblies through all brain states, and liquidity was brain region- and brain state-specific (Figure 4C). As reported in Table 2, the largest liquidity was observed for mPFC sharing assemblies during natural sleep (∼94%). CA1 displayed a substantial reduction in the liquidity of sharing assemblies when moving from anesthesia to sleep (dropping from ∼86% in anesthesia to ∼57% in sleep). Finally, as for the other features, information sharing substates were brain state specific (Figure 4D).

**Table 2.** Sharing assembly liquidity across regions and conditions

| Region and state | Liquidity sharing strengths | Liquidity sharing assemblies |
| --- | --- | --- |
| mEC (*anesthesia*) | 0.31 (0.22, 0.45) | 0.84 (0.72, 0.89) |
| CA1 (*anesthesia*) | 0.39 (0.36, 0.47) | 0.86 (0.77, 0.89) |
| CA1 (*sleep*) | 0.04 (0.03, 0.05) | 0.57 (0.50, 0.68) |
| mPFC (*sleep*) | 0.18 (0.11, 0.26) | 0.94 (0.89, 0.96) |

*Median (1<sup>st</sup>, 3<sup>rd</sup> quartile), SO/THE and non-REM/REM states confounded*

### Loose coordination of substate transitions between brain regions

Single units were recorded simultaneously in two regions (CA1 and mEC; CA1 and mPFC). We thus assessed whether substate transition events in one region matched the transition in the other region. We computed the relative mutual information between substate sequences of a given type (e.g. firing, storage or sharing) observed in one region and the other. We did not find significant differences for these measures across the three features (firing, storage, sharing) and therefore pooled them together. The median relative mutual information between substate transitions in the probed cortical and hippocampal regions was 18% during anesthesia (between mEC and CA1) and 42% during natural sleep (between mPFC and CA1). These levels of coordination between substate sequences denoted a lack of perfect parallelism between transitions in the different regions, but they were still well above chance level (Figure S10). Thus, substate dynamics display some coordination between CA1 and mPFC during sleep (Table 3), which is in keeping with the fact that information exchange occurs between the two regions during sleep (*15*). The weak coordination under anesthesia suggests that circuits may operate more independently from one another in this condition (but still not completely).

**Table 3.**
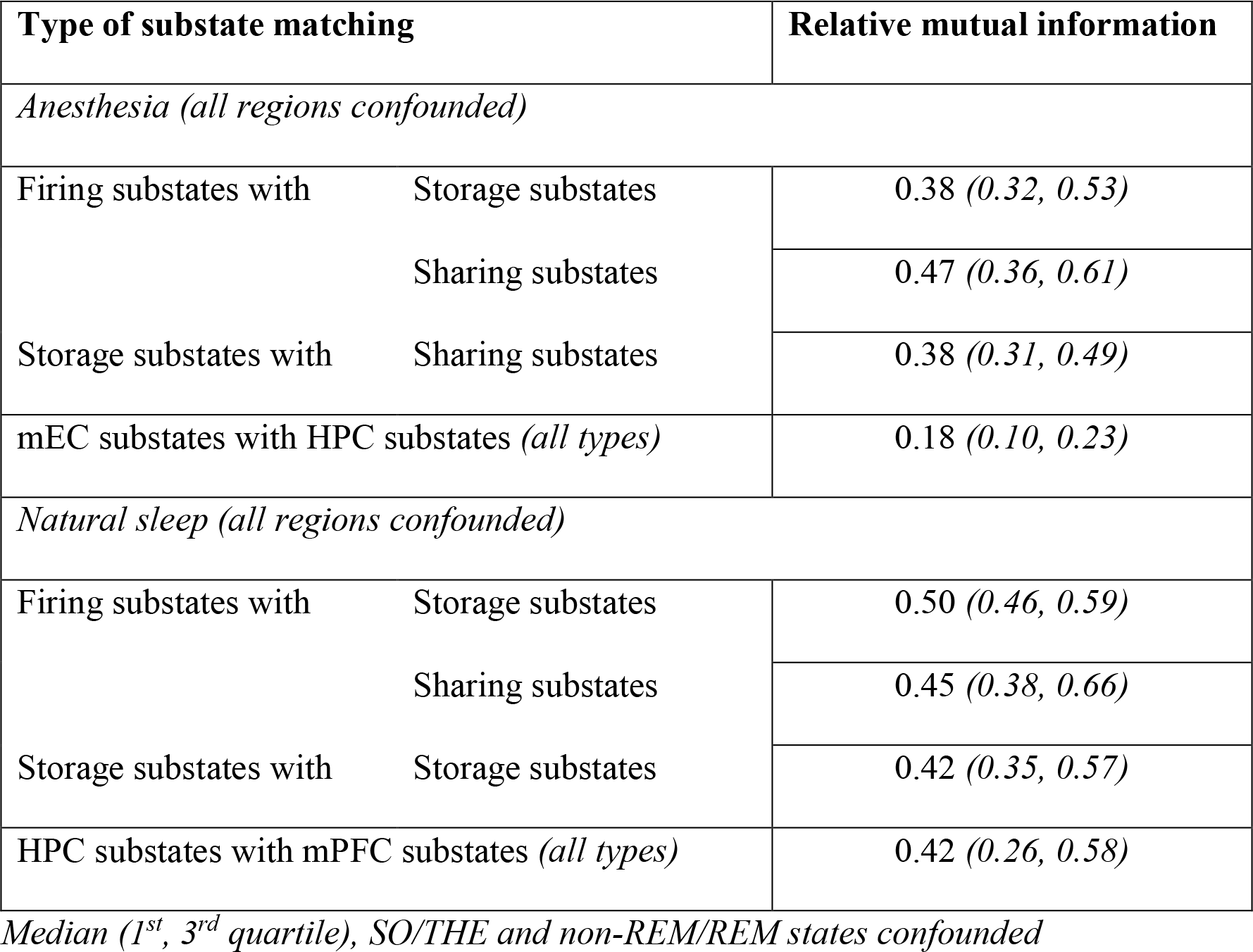
Matching between substate sequences of different types across conditions and regions

### A large fraction of cells can act as computing hubs

Functional, effective, and anatomical hub neurons (mostly GABAergic) have been identified in the brain (*16*). We complement the concept, introducing *storage* and *sharing hubs*, i.e. neurons displaying an elevated storage or sharing values, respectively (see Methods). In contrast to the sparsity of functional, effective, and anatomical hubs, a large fraction of cells acted as a computing hub in at least one substate, as illustrated in Figure 5A. Computing hubs could be recruited across all probed regions and layer locations (Figure 5B and C). As summarized in Figure 5B, the probability of serving as computing hub – storage or sharing confounded – was of 40% or more on average for almost all layers, apart from the, possibly under-sampled, stratum lacunosum moleculare and stratum radiatum in CA1. We observed a general tendency for inhibitory interneurons to have a larger probability to serve as computing hubs than for excitatory cells. This tendency was particularly strong for cortical regions and was notably significant in layer 5 of mEC (during anesthesia) and layer 3 of mPFC (during sleep), for which the probabilities of inhibitory interneurons serving as computing hub in at least one substate approached 70%. The probability of serving as a computing hub at least once was relatively similar when evaluated separately for storage or sharing. In particular, 43% of the neurons serving as a storage hub in a substate could serve as a sharing hub in another substate, but in general, not simultaneously as only 12% of the neurons were “multi-function” hubs.

**Figure 5.**
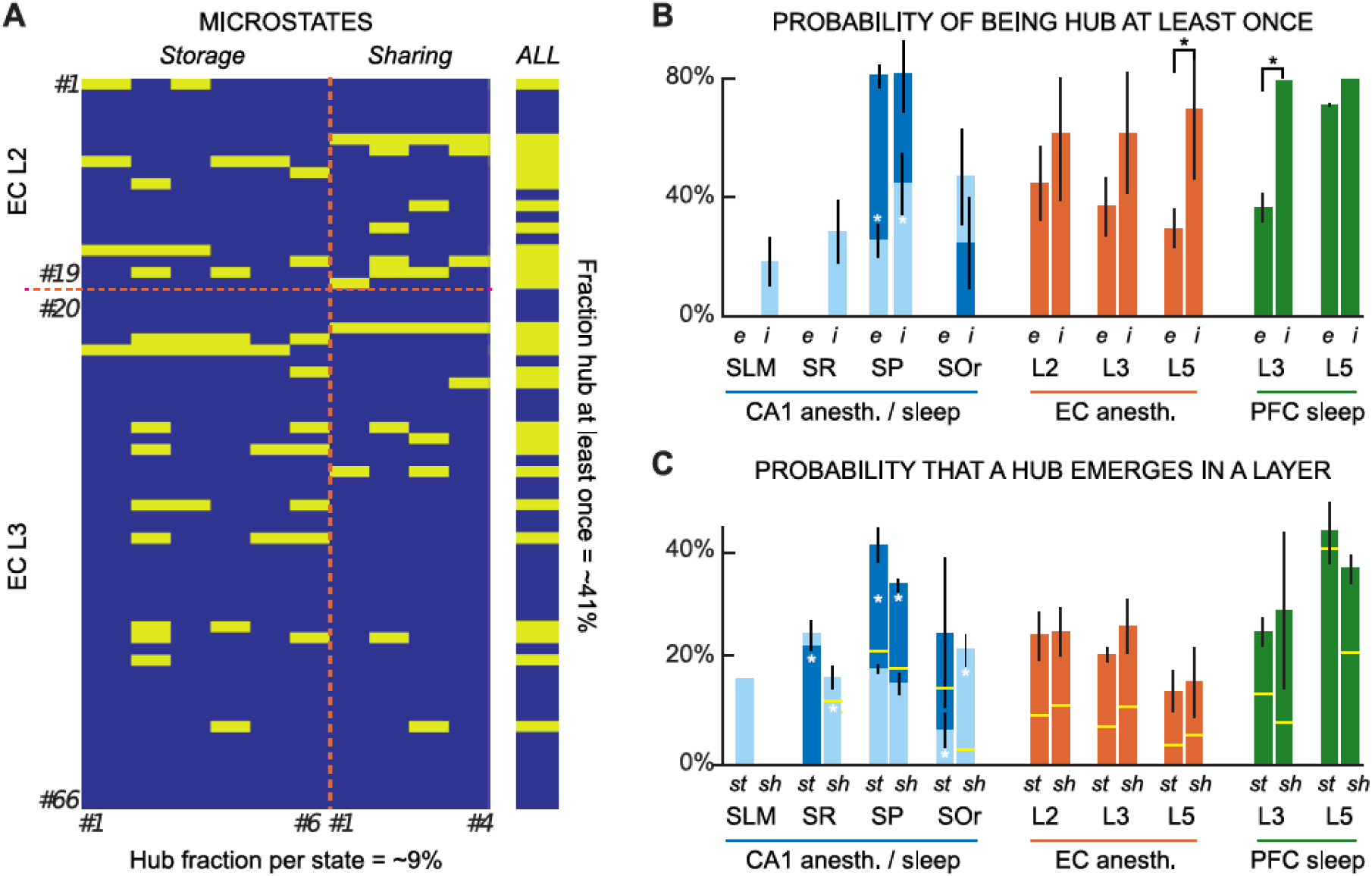
A democracy of computing hubs. **(A)** Within every computing substate some neurons exhibited significantly strong values of information storage or sharing (*computing hubs*). However, these computing hubs did generally change from one substate to the other, as shown in this example. Different rows correspond to different single units recorded in mEC during anesthesia and different columns correspond to different computing substates (left, storage substates 1 to 6; right, sharing substates 1 to 4). An entry is colored in yellow when the neuron is a computing hub within the corresponding substate. In the shown example, while ∼9% of neurons on average were simultaneously acting as computing hub, over 40% of the recorded units were recruited as hubs for at least one substate, when considering all the computing substates together (vertical bar on the right). **(B-C)** The probability that a neuron acted as hub depended only loosely on its anatomical localization. Panel **B** shows that for all regions and layers the probability that a neuron act as computing hub at least once was always larger than 30%. Inhibitory (*i*) neurons tended to be recruited as hubs more frequently than excitatory (*e*) neurons. Analogously, panel **C** shows that none of the layers display a specialization in either one of the two processing operations of information storage or sharing. Stars denote statistically significant comparisons (lack of overlap between 95% confidence intervals for the probability, reported as vertical ranges on top of the histogram bar). In panel **C** a yellow horizontal line indicates the fraction of computing hub cells which also happen to simultaneously be high-firing rate cells. Many computing hubs have thus average or low firing rate. In panel **B-C** in CA1 light blue represents anesthesia and dark blue represents natural sleep.

Despite this large flexibility in the dynamic assignment of hub roles, the notion of hub continued to make sense within each individual substate. Within a substate, on average only ∼9% of cells acted as hub (storage or sharing pooled), so still a strict “elite” (although not a permanent one but appointed just within the associated state). The set of recruited hubs constituted thus at each time a characteristic fingerprint of the active substates (with only 4% of the substates being “hubless”).

We also studied the probability that a computing hub emerged in a given layer (Figure 5C). During anesthesia, all probed layers of CA1 and mEC showed a ∼20% uniform probability for a storage and sharing computing hub to emerge. Natural sleep was associated to an enhanced recruitment of computing hubs. The probabilities of hub emergence exceeded ∼40% for storage hubs in layer 5 of mPFC and in SP of CA1. The analysis of deep or superficial CA1 SP principal neurons, which are involved in different microcircuits (*17*-*18*), did not reveal an intra-layer distribution of computing hubs (not shown). These results suggest that the probability that a neuron serves as computing hub is not correlated to its anatomical region or layer location.

Finally, we tested whether computing hubs were characterized by high firing rates. Using the same procedure utilized to extract computing hubs, we found that 62% of the cells were high-firing at least in one firing substate with 70% being putative interneurons. Remarkably, there was a poor overlap between computing hubs and high firing rate cells. Table 3 already shows that storage and sharing substate sequences are only loosely coordinated with firing substate sequences (cf. as well firing rate information in Figures 3B-E and S8B-E). Furthermore, being a high firing rate cell does not guarantee that this cell will also be a computational hub (or the other way around). This is also shown in Figure 5C, where the yellow levels over the histogram bars indicate the fraction of storage and sharing hubs which also happen to be high firing cells. We conclude that a storage hub can have a normal or even smaller than average firing rate.

### The syntax of substate sequences is complex and brain state-dependent

Collectively, our results demonstrate the existence of substate sequences in three different brain regions during anesthesia and natural sleep. Using a linguistics analogy (Figure S11A), we assign a *letter* to each identified substate (represented by a color in the figures). The temporal sequence of substates thus translates into a stream of *letters*. However, if we consider the three features simultaneously, we obtain a stream of 3 letter *words* (Figure S11B). All combinations of possible letters from our 3 features define the *dictionary* of *words* that can be expressed. We represent a stream of *words* as a switching table (Figure 6A). This allows us to explore two aspects of the “neuronal language”: the statistics of the *words* and the statistics of the transitions between the words (the *syntax*). We found that the *words* were mostly (85%) brain state-specific, as expected since the substates *letters* are already brain state-specific (cf. Figures 3C and F, 4D). Although the syntactic rules structuring the production of *words* are unknown, we can quantify their complexity. Algorithmic information theory (*19*), the minimum description length framework (*20*) and the Lempel-Ziv method (*21*) link complexity to the notion of compressibility. As illustrated in Figure 6B, an ordered, regular switching table requires a short description, as a small list of instructions can be written to reproduce the table (e.g. *word D* 100 times, followed by *word B* 88 times, etc.). At the opposite extreme, a completely random switching table would need a lengthy exhaustive description -as many instructions as the length of the table itself. A complex switching table stands between regularity and randomness and requires a description that is compressed, longer than a regular table but shorter than a random table.

**Figure 6.**
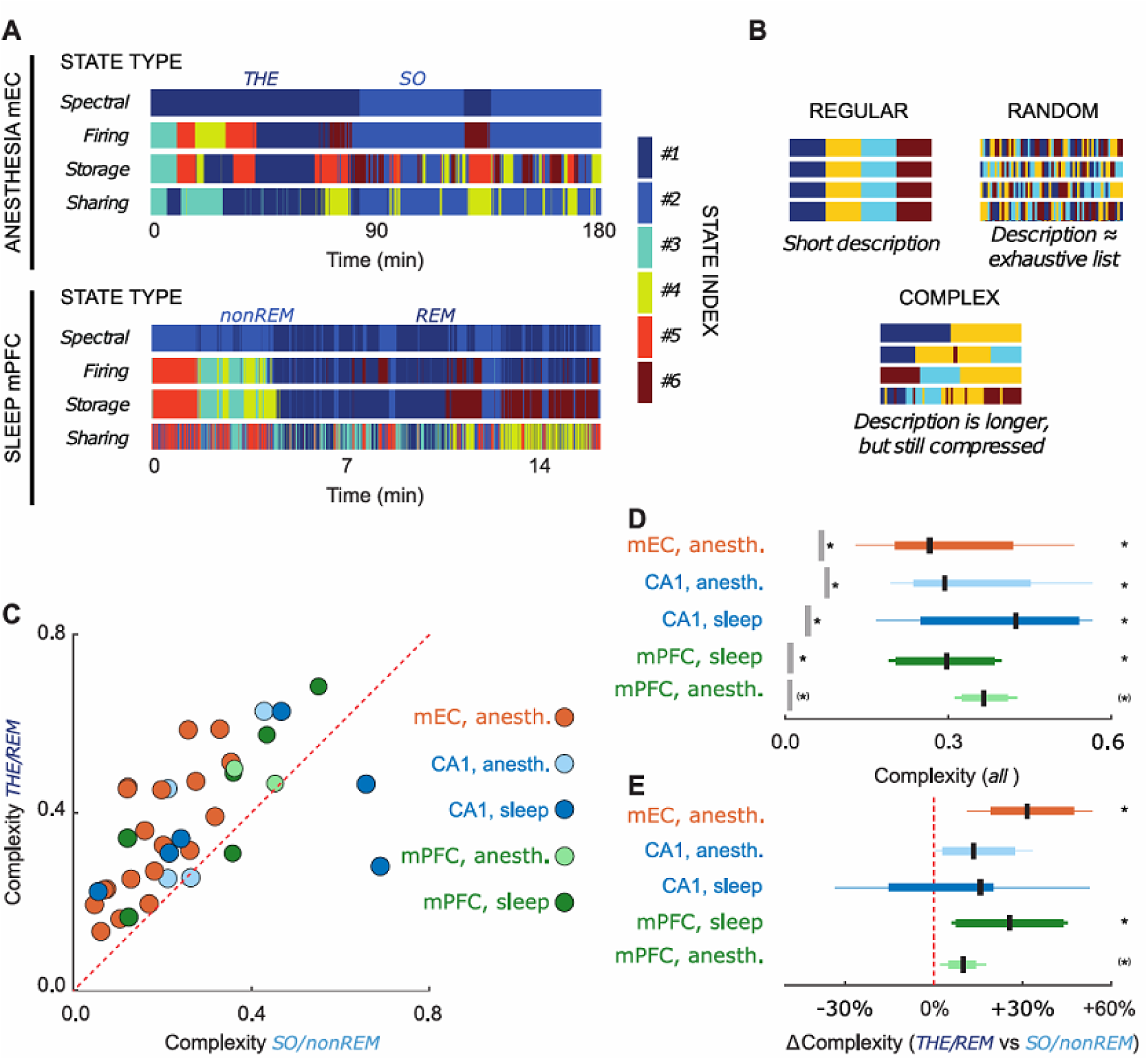
Complexity of substate sequences. State switching found for each feature (firing, storage, sharing) did not align in time. This can be visualized by state switching tables, whose different rows graphically represent transitions between global brain oscillatory states and firing, storage, and sharing substates. Examples of switching tables are shown in **(A)** for mEC during anesthesia (top) and for mPFC during natural sleep (bottom, note the different time scales). Switching tables were neither perfectly regular (**B**, top left), nor random (**B**, top right), but they were “complex”, displaying organized patterns escaping simple description (**B**, bottom). **(C)** The complexity of the switching tables was larger for THE/REM than for SO/nonREM for most recordings. We included two recordings from mPFC under anesthesia for comparison. (**D**) Switching tables were complex in all cases. Complexity values were significantly above the upper threshold for regularity and below the lower threshold for randomness. (**E**) The increase of complexity was significant for mEC when transitioning from SO to THE and for mPFC from nonREM to REM sleep. This trend in CA1 was not statistically significant (significance assessed in terms of lack of intersection between 95%-confidence intervals and threshold values for both panels **D** and **E**; a (*) symbol indicates that the number of recordings in this category was not enough to assess significance but that the median value lied below or above the considered threshold).

Figure 6C shows that the syntax was complex (between 0 and 1) for all brain regions and brain states and that THE/REM states were more complex than SO/nonREM states. We added two recordings from mPFC under anesthesia for comparison. Figure 6D shows that the measured complexity was significantly larger than the upper threshold for regularity and significantly smaller than the lower threshold for randomness (p<0.05, Bonferroni Corrected, direct c.i. comparison).

Finally, we assessed whether switching from SO to THE or from nonREM to REM increased the complexity. As shown in Figure 6E, the tendency was toward an increase of complexity in all cases, from +30% for mEC during anesthesia and mPFC during anesthesia or sleep to roughly +10% for CA1 during anesthesia or sleep. This relative increase was always significant (*p* < 0.05, Bonferroni Corrected, c.i. comparison) apart from CA1, for which two recordings displayed increased complexity during nonREM sleep. We conclude that the syntax is complex and brain state-dependent.

### What determines complexity?

We then investigated which factors contribute to complexity. Different durations of *words* may account for variations in complexity. Although *word* dwell times were different by one order of magnitude between anesthesia and sleep with median ∼18 min (∼10 min 1^st^ quartile, ∼28 min 3^rd^ quartile) during anesthesia and ∼1.4 min (∼1 min 1^st^ quartile, ∼2.1 min 3^rd^ quartile) during sleep, complexity values for anesthesia and natural sleep were similar.

We also evaluated the burstiness coefficient, *B* (*22*), of the stream of *words*. This coefficient ranged between −1 ≤ *B* ≤ 1, with *B* = −1 corresponding to a perfectly periodic stream of *words*, *B* = 0 to a Poisson train and *B* = 1 to a maximally bursting stream. We found a positive correlation between burstiness and complexity (Figure S11A, *p*<0.01, Bootstrap c.i). Burstiness was greater during THE/REM (0.15) than during SO/nonREM (0.09 *p* = 0.03, Kruskal-Wallis test), which may contribute to the increased complexity found during THE/REM.

The richness of the dictionary also affects complexity (*21*). We therefore evaluated the Used Dictionary Fraction, i.e. the ratio between the number of observed *words* and the maximum theoretical number of *words*, i.e. the *dictionary*. We find a significant positive correlation between the Used Dictionary Fraction and complexity (Figure S12B, *p*<0.05, Bootstrap c.i). The richness of the dictionary was greater during THE/REM (21%) than during SO/nonREM (14%, *p* = 0.032, Kruskal-Wallis test), which may also contribute to the increased complexity found during THE/REM.

A bivariate linear regression of complexity over burstiness and Used Dictionary Fraction revealed a correlation of 0.62 (*p*<0.05, Bootstrap c.i) between predicted and observed complexity, demonstrating that complexity is largely explained by burstiness and the Used Dictionary Fraction.

Finally, we verified that our results did not depend on the measure of complexity. Redoing analyses using Lempel-Ziv complexity (*21*), which was previously used to analyze neural activity (*23*-*24*), lead to qualitatively equivalent results. Lempel-Ziv complexity also strongly correlated with our measure of complexity (Figure S11C Pearson correlation 0.84, *p* < 0.001, bootstrap c.i.).

## Discussion

Here we demonstrate two levels of organization of brain activity. At the single cell level, we find that a large proportion of recorded neurons act as computing hubs during discrete time epochs (substates) within a given stable brain state (e.g. REM and nonREM). At the microcircuit level, we find a rich repertoire of computational substates characterized by temporally structured sequences, whose complexity was modulated by the global brain oscillatory state. Such type of organization was shared between three anatomical different brain regions: the hippocampus, the medial entorhinal cortex and the medial prefrontal cortex.

The “hubness” of a neuron may be determined by fixed features, e.g. an exceptional extension of axonal arborizations (*25*); a suitable location in the circuit wiring diagram facilitating the control of synchronization (*26*); or yet, some developmental “droit d’aînesse” (*16*). During natural sleep and anesthesia, however, we find that >40% of the recorded neurons act as a computational hub during at least one substate, meaning that computing hubs form a rather open and not so elitist club. The computational hubness is dynamic - a neuron acting as a hub in a given substate may not be a hub in a different substate or may swap its nature (e.g. converting from a storage to a sharing hub). The stronger tendency for putative inhibitory cells to serve as hubs (>70%) is in keeping with the known role of GABAergic cells in orchestrating network activity (*16*, *25*-*26*). Furthermore, because our analysis was limited to few brain states, the proportion of putative principal and GABA neurons acting as computational hubs may even be an underestimate. Perhaps all neurons act as computational hubs during specific brain states (including exploration and quiet awakening).

That hubs share information with ever changing source and target neurons is in apparent contradiction with the existence of sequential firing of cell assemblies in cortical and hippocampal circuits, including during nonREM sleep (*11*,*16*, *27*-*37*). Our information-theoretical analyses require the use of at least 5 s long sliding windows, which is not sufficient to detect fast sequences of activation, as replay events occur within 500 ms (*38*). Interestingly, replay sequences are not strictly stable as they demonstrate inter-cycle variability (*39*), which may reflect liquidity. The liquid nature of information sharing suggests that neuronal activity is not frozen at fixed-point attractors as in classic artificial neural networks (*40*) but may be sampling the broad basin of attraction of shallow attractors (*8*) or higher-dimensional manifolds “at the edge of chaos”, as found in reservoir computing schemes (*41*-*43*). In this case, information is shared across extremely volatile assemblies within a given substate. The assembly dynamics are thus “liquid” – i.e. neither frozen into crystallized patterns, nor fully random as in a gas – and are only mildly constrained to robustly maintain the computational role of sharing hubs while preserving entropy of firing patterns, and therefore bandwidth for local computations (*44*). This preservation of hub function in a heterogeneous and reconfiguring circuit can be seen as a form of homeostasis of the computing role, generalizing the concept of homeostasis of circuit behavior evidenced in invertebrate systems (*45*-*46*). While this latter homeostasis preserves the functional level, in our case homeostasis would extend down to the algorithmic level, referring to the three-level hierarchy proposed by Marr & Poggio (*1*).

During a “stable” behavior such as resting state, analysis of BOLD and EEG signals consistently revealed the presence of temporal sequences of resting state networks and topographical microstates, respectively (*4*-*5*). Here, we find that an analogous switching between discrete states occurs at a completely different scale of microcircuits. During a “stable” oscillatory regime (e.g. theta rhythm), neuronal computation is indeed organized in temporal sequences of computational substates. Interestingly, while field oscillations constrain neuronal firing and neuronal firing produces field oscillations (*47*), we find only a loose match between the switch from one oscillatory mode to the other and the switch from one substate to the other. Transitions between global states – related to the scale of mesoscale collective dynamics– sometimes anticipate and sometimes follow transitions between firing, storage or sharing substates –related to the scale of microscopic firing dynamics–, as if dynamic changes occurring at either one of the scales had destabilizing effects on the dynamics at the other scale (in both directions, meso- to micro-scale and micro-to meso-scale). The behavior of CA1 cells may reflect specific internal dynamics, not tightly controlled by the CA1 local field which mostly reflects synaptic inputs originating from outside the CA1 region. Importantly, the repertoire of computing substates is brain state specific. Beyond proposals that oscillations are central for the routing of information between regions (*47*-*48*), we thus suggest here that global oscillatory states could also organize information processing within local regions by enforcing the use of their own state-specific “languages” (expressed in terms of combinations of alternative intrinsic substates).

Signatures of computation can be identified even if the function and meaning of the computation are unknown and even when system states are sampled partially, as it is the case for the present study. This allowed us to extract a symbolic representation of substates (*letters*) for a given feature, which make *words* when considering several features. The syntax of the substate word sequences is complex, standing between order and randomness (as it was already the case for the sharing dynamics within each substate). The capacity to generate complex sequences of patterns is a hallmark of self-organizing systems and has been associated to their emergent potential to perform universal computations (70). Moreover, dynamics at the “edge of chaos” confer advantages for information processing (41–43).

Importantly, we find that the syntactic complexity of substate sequences is brain state-dependent as it was the case for the substate dictionaries, and more complex during theta oscillations/REM sleep than during slow oscillations/nonREM sleep, suggesting an increased load of computation in the former brain state. Remarkably, the temporal complexity of activation sequences was also shown to be modulated by brain states at the macro-scale level of whole-brain dynamics (*49*). In keeping with the view that slow/theta oscillations measured during anesthesia share general properties with nonREM/REM sleep (*50*-*52*), we found similar rules of organization in terms of substate sequences and their complexity, despite the fact that the *word* dwell times in anesthesia are one order of magnitude greater than during natural sleep. We speculate that the nature of the undergoing oscillation (slow vs theta) constrains the repertoire of *words* used and their syntax, modulating the type of computation performed by the recruitment of varying computing hubs. Sleep, oscillatory patterns, and neuronal firing are altered in numerous neurological disorders, including epilepsy (*18*, *53*-*56*) and therefore it will be important to assess whether the repertoire of substates and the syntax are likewise affected.

In conclusion, our results reveal a rich algorithmic-level organization of brain computations during natural sleep and anesthesia, which combines a complex combinatorics of discrete states with the flexibility provided by liquidly reconfiguring assemblies. While we cannot yet prove that this substate dynamics is functionally relevant, it has the potential to serve as a substrate for previously undisclosed computations. The next aim will be to perform the similar analysis during specific behavioral tasks, such as goal-driven maze navigation. *Words* and/or their sequence may sign specific cognitive processes. The fact that the algorithmic instructions and primitive processing operations are similar in three brain regions with different architectural organizations suggests the existence of a basic architecture for low-level computations shared by diverse neuronal circuits.

## Acknowledgements

We used part of the data collected in G. Buzsáki’s laboratory and originally published by Quilichini et al. (2010) for our investigations. We acknowledge G. Buzsáki and A. Sirota for consenting to the use of this database in the present work. We also used part of the data originally published by Ferraris et al. (2018). PPQ acknowledges support from FRM, FFRE and CURE Epilepsy Taking Flight Award. MF acknowledges support from FRM. The M-GATE project has received funding from the European Union’s Horizon 2020 research and innovation program under the Marie Skłodowska-Curie grant agreement No 765549. DB acknowledges support by the CNRS “Mission pour l’Interdisciplinarité” INFINITI program (BrainTime) and the French Agence Nationale pour la Recherche (ERMUNDY, ANR-18-CE37-0014-02). The funders had no role in study design, data collection and analysis, decision to publish, or preparation of the manuscript. We wish also to acknowledge Viktor Jirsa, Lionel Barnett and Thilo Womelsdorf for constructive comments.

## Materials and Methods

### Data information

We use in this work a portion of the data (13 out of 18 experiments) initially published by Quilichini et al. (2010), which includes local field potentials (LFPs) and single-unit recordings obtained from the dorsomedial entorhinal cortex (mEC) of anesthetized rats. We also use a portion of the data (2 out of 16 experiments) initially published by Ferraris et al. (2018), which includes LFPs and single-units recorded in the medial prefrontal cortex (mPFC) under anesthesia. Seven recordings are original data in both mEC and dorsal hippocampus (HPC) under anesthesia, and 10 recordings in 4 animals during natural sleep in HPC and mPFC. See Figures S1 and S2 for details on recordings, number of cells, and layers recorded.

### Animal surgery

We performed all experiments in accordance with experimental guidelines approved by the Rutgers University and Aix-Marseille University Animal Care and Use Committee. We performed experiments on 13 male Sprague Dawley rats (250–400 g; Hilltop Laboratory Animals), 8 male Wistar Han IGS rats (250-400g; Charles Rivers) and 3 male Long Evans rats (350-400g; Charles River). We performed acute (anesthesia) experiments on the Sprague Dawley and 7 of the Wistar rats, which were anesthetized with urethane (1.5 g/kg, i.p.) and ketamine/xylazine (20 and 2 mg/kg, i.m.), additional doses of ketamine/xylazine (2 and 0.2 mg/kg) being supplemented during the electrophysiological recordings. We performed chronic (natural sleep) experiments on one Wistar and the Long Evans rats, which were anesthetized using isoflurane 2% in 1l/min of O_2_ for the surgery procedure. In both cases, the body temperature was monitored and kept constant with a heating pad. The head was secured in a stereotaxic frame (Kopf) and the skull was exposed and cleaned. Two miniature stainless-steel screws, driven into the skull, served as ground and reference electrodes. To reach the mEC, we performed one craniotomy from bregma: −7.0 mm AP and +4.0 mm ML; to reach the CA1 area of HPC, we performed one craniotomy from bregma: − 3.0 mm AP and +2.5 mm ML in the case of HPC coupled to mEC recordings, and from bregma: −5.6 mm AP and +4.3 mm ML-3.0 mm in the case of HPC coupled to mPFC recordings; to reach the mPFC, we performed one craniotomy from bregma: +3 mm AP and +0.8 mm ML. We chose these coordinates to respect known anatomical and functional connectivity in the cortico-hippocampal circuitry (*51, 57-59*). We used different types of silicon probes to record the extracellular signals. In acute experiments, the probes were mounted on a stereotaxic arm. We recorded the dorso-medial portion of the mEC activity using a NeuroNexus CM32-4×8-5mm-Buzsaki32-200-177 probe (in 8 experiments), a 10-mm long Acreo single-shank silicon probe with 32 sites (50 µm spacing) arranged linearly (in 5 experiments), or a NeuroNexus H32-10mm-50-177 probe (in 5 experiments), which was lowered in of the EC at 5.0-5.2 mm from the brain surface with a 20° angle. We recorded HPC CA1 activity using a H32-4×8-5mm-50-200-177 probe (NeuroNexus Technologies) lowered at 2.5 mm from the brain surface with a 20° angle (in 4 experiments), a NeuroNexus H16-6mm-50-177 probe lowered at 2.5 mm from the brain surface with a 20° angle (in 2 experiments) and a E32-1shank- 40µm-177 probe (Cambridge Neurotech) lowered at 2.5 mm from the brain surface with a 20° angle (in 1 experiment). We recorded mPFC activity using NeuroNexus H32-6mm-50-177 lowered in the layer 5 at 3 mm perpendicularly from the brain surface (in 2 experiments). In chronic experiments, the probes were mounted on a movable micro-drive (Cambridge Neurotech) fixed on the skull and secured in a copper-mesh hat. We recorded HPC CA1 activity (probes lowered perpendicularly at 2.5 mm from the brain surface) using a Neuronexus H32-Poly2-5mm-50-177 probe (in 2 experiments), a Cambridge Neurotech E32-2shanks-40µm-177 probe (in 1 experiment) and a NeuroNexus H32-4×8-5mm-50-200-177 probe (in 1 experiment). We recorded mPFC activity (probes lowered perpendicularly at 3.0 mm from the brain surface) using a NeuroNexus H32-4×8-5mm-50-200-177 probe (in 2 experiments), and a Neuronexus H32-Poly2-5mm-50-177 probe (in 1 experiment). The on-line positioning of the probes was assisted by: the presence of unit activity in cell body layers and the reversal of theta ([3 6] Hz in anesthesia, [6 11] Hz in natural sleep) oscillations when passing from L2 to L1 for the mEC probe, and the presence in stratum pyramidale either of unit activity and ripples (80-150 Hz) for the HPC probe, and the DV depth value and the presence of intense unit activity for the mPFC.

At the end of the recording, the animals were injected with a lethal dose of Pentobarbital Na (150mk/kg, i.p.) and perfused intracardially with 4% paraformaldehyde solution. We confirmed the position of the electrodes (DiI was applied on the back of the probe before insertion) histologically on Nissl-stained 40 µm section as reported previously in detail (*60*). We used only experiments with appropriate position of the probe for analysis. The numbers of recorded single units in different anatomical locations for the different retained recordings are summarized in Figure S2.

### Data collection and spike sorting

Extracellular signal recorded from the silicon probes was amplified (1000x), bandpass filtered (1 Hz to 5 kHz) and acquired continuously at 20 kHz with a 64-channel DataMax System; RC Electronics or a 258-channel Amplipex, or at 32 kHz with a 64-channel DigitalLynx; NeuraLynx at 16-bit resolution. We preprocessed raw data using a custom-developed suite of programs (*61*). After recording, the signals were downsampled to 1250 Hz for the local field potential (LFP) analysis. Spike sorting was performed automatically, using KLUSTAKWIK (http://klustakwik.sourceforge.net (*62*)), followed by manual adjustment of the clusters, with the help of auto-correlogram, cross-correlogram and spike waveform similarity matrix (KLUSTERS software package, http://klusters.source-forge.net (*63*)). After spike sorting, we plotted the spike features of units as a function of time, and we discarded the units with signs of significant drift over the period of recording.

Moreover, we included in the analyses only units with clear refractory periods and well-defined clusters. Recording sessions were divided into brain states of theta and slow oscillation periods. The epochs of stable theta (THE in anesthesia experiments, REM in natural sleep experiments or slow oscillations (SO in anesthesia experiments, non-REM in natural sleep experiments) periods were visually selected from the ratios of the whitened power in the theta band ([3 6] Hz in anesthesia, [6 11] Hz in natural sleep) and the power of the neighboring bands ([1 3] Hz and [7 14] Hz in anesthesia, [12 20] Hz in natural sleep) of EC layer 3 LFP, which was a layer present in all the 18 anesthesia recordings, or layer 5 mPFC recordings in natural sleep recordings, and assisted by visual inspection of the raw traces (*60*) (Figure S3). We then used band-averaged powers over the same frequency ranges of interest as features for the automated extraction of spectral states via unsupervised clustering, which confirmed our manual classification.

We determined the layer assignment of the neurons from the approximate location of their somata relative to the recording sites (with the largest-amplitude unit corresponding to the putative location of the soma), the known distances between the recording sites, and the histological reconstruction of the recording electrode tracks.

### Characterizations of single unit activity

We calculated pairwise cross-correlograms (CCGs) between spike trains of these cells during each brain state separately (60, 64-65). We determined the statistical significance of putative inhibition or excitation (trough or peak in the [+2 5] ms interval, respectively) using the nonparametric test and criterion used for identifying monosynaptic excitations or inhibitions (60, 64-65), in which each spike of each neuron was jittered randomly and independently on a uniform interval of [-5 5] ms a 1000 times to form 1000 surrogate data sets and from which the global maximum and minimum bands at 99% acceptance levels were constructed. Inspection of CCGs thus allowed to identify single units as putatively excitatory or inhibitory, an information which we used to perform the computing hub characterizations in Figure 5B.

To perform the analyses of Figure S4 we then computed the burst index and the phase modulation of units. Burst index denotes the propensity of neurons to discharge in bursts. We estimated the amplitude of the burst spike auto-correlogram (1 ms bin size) by subtracting the mean value between 40 and 50 ms (baseline) from the peak measured between 0 and 10 ms. Positive burst amplitudes were normalized to the peak and negative amplitudes were normalized to the baseline to obtain indexes ranging from −1 to 1. Neurons displaying a value of 0.6 were considered bursting.

To establish the phase modulation of units, we concatenated different epochs of slow or theta oscillations, and estimated the instantaneous phase of the ongoing oscillation by Hilbert transform of the [0.5 2] Hz or [3 6] Hz in anesthesia and [6 11] Hz in natural sleep filtered signal, for slow or theta oscillations respectively. Using linear interpolation, we assigned a value of phase to each action potential. We determined the modulation of unit firing by Rayleigh circular statistics; *p* < 0.05 was considered significant. We first assessed circular uniformity of the data with a test for symmetry around the median (*66*) and we performed group comparison tests of circular variables using circular ANOVA for uniformly distributed data and using a nonparametric multi-sample test for equal medians “CM-test”, similar to a Kruskal–Wallis test, for non-uniformly distributed data (Berens, 2009; https://philippberens.wordpress.com/code/circstats), and p < 0.05 was considered significant.

### Feature-based state extraction

We performed a sliding-window analysis of the recorded LFP time-series and single unit spike trains, extracting in a time-resolved manner a variety of different descriptive features. For all the considered features (see specific descriptions in later sub-sections), we use similar window sizes and overlap for the sake of a better comparison. For anesthesia recordings, we adopted a long window duration of 10 s – demanded by the estimation needs for the most “data-hungry” information-theoretical features – with an overlap of 9 s. For natural sleep recordings, we adopted a window duration of 10 s with an overlap of 9 s.

We computed each set of descriptive features and compiled them into multi-entry vectors `FeatureVector`*(t)* for every time-window centered on different times *t*.

We then compute a similarity matrix M_sim_, to visualize the variability over time of the probed feature set. The entries *M_sim_(t_a_, t_b_)* are given by the Pearson correlation coefficient between the entries in the vectors `FeatureVector`*(t_a_)* and `FeatureVector`*(t_b_)*, treated as ordered sequences, and are thus bounded between −1 and +1. Blocks of internally elevated correlation along the similarity matrix diagonal denote epochs of stable feature configurations. Similar configurations are detected by the presence of off-diagonal highly-internally correlated blocks and the existence of multiple possible configurations by the poor correlation between distinct blocks.

We then extracted feature-based states using a standard iterative *K*-means algorithm (*67*) to cluster the different vectors `FeatureVector`*(t),* based on the correlation distance matrix defined by 1-*M_sim_*. We defined the substates of different types as the different clusters obtained for different feature types. We chose the number of clusters K by clustering using *K* = 2, 3,… 20 and first guessing *K* using a maximal silhouette criterion (*68*) across all *K*s. We also inspected dendrograms from single-linkage clustering as a cross-criterion. Using both pieces of information the *K* was manually adjusted case-by-case (up to ±2 clusters with respect to the unsupervised silhouette criterion) to best match the visually apparent block structure of the similarity matrix *M_sim_*, which results in an optimized K selection for each recording.

### Feature Robustness

To compute the robustness of the feature computation, the original spiking times were randomly shuffled 1000 times and the features recomputed for each instance for 2 files, one in anesthesia and one in natural sleep. To compare it to the original features computed, the *k* for each recording and each feature was kept the same. The information retained after shuffling was computed by dividing the mutual information between the shuffled features and the original by the entropy of the new feature set. We found a significant difference for both anesthesia and in natural sleep across all features, and the results have been quantified in Table S1.

### Global oscillatory states

We defined eight different unequally-sized frequency ranges, which were manually adjusted recording-by-recording to be better centered on the recording-specific positions of the slow-wave and theta peaks and of their harmonics (e.g., 0–1.5 Hz, 1.5–2 Hz, 2–3 Hz, 3–5 Hz, 5–7 Hz, 7–10 Hz, 10–23 Hz and 23–50 Hz for the anesthesia spectrogram and the similarity matrix of Figure S3A). We averaged the spectrograms over all channels within each of the layers in the simultaneously recorded regions (e.g. EC and CA1 for anesthesia) and then we coarse-grained the frequencies by further averaging over the eight above ranges. We compiled finally all these layer-averaged and band-averaged power values into time-dependent vectors `Spectra`*(t)*, with a number of entries given by eight (number of frequency bands) times the number of layers probed in the considered recording, i.e. up to eight (CA1 stratum oriens (SOr), stratum pyramidale (SP), stratum radiatum (SR) and stratum lacunosum moleculare (SLM); EC layers 2, 3 and 5; and PFC layers 1,2, 3 and 5), yielding at most 64 entries. We then processed these spectral features as described in the previous section to extract global oscillatory states –as any other substate type– via unsupervised clustering.

### Firing sets and firing hubs

Not all neurons are equally active in all temporal windows. To determine typical patterns of single neuron activation we binned the spiking data for each unit in 50 *ms* windows – if a neuron fired within that window the result was a ‘1’, if it did not fire the result was a ‘0’. We enforced a strictly binary encoding, i.e. we attributed to a bin a ‘1’ symbol even when more than one spike was fired within this bin. Our bin size choice however was such to maintain the loss of information when ignoring multiple firing events within a bin was less that 5%. Note furthermore that for the majority of spike trains multiple firing events were extremely rare, i.e. apart from a few cases the information loss was way smaller than 5%. We then averaged over time this binned spike density, separately for each single unit and within each time window and compiled these averages into time-dependent vectors `Firing`*(t)*, with *N* entries, where *N* is the overall number of single units probed within the considered recording. We constructed separate feature vectors for each of the simultaneously recorded regions. Firing substate prototypes were given by the centroids of the clusters extracted from the similarity matrix *M_sim_* resulting from the stream of `Firing`*(t)* feature vectors.

We then defined a neuron to be a *high-firing cell* in a given state if its firing rate in the state prototype vector was higher than the 95% percentile of all concatenated state prototype vector entries.

### Active Information Storage

Within each time-window we computed for each single unit an approximation to the Active Information Storage (AIS). AIS is meant to quantify how much the activity of a unit is maintaining over time information that it was conveying already in the past (*3*, *6*). This information-theoretical notion of storage is distinct from the neurobiological notion of storage in synaptic weights. It is indeed an activity-based metric (hence the adjective “active”), able to detect when temporal patterns in the activity of a single unit can serve the functional role of “memory buffer”. AIS is strictly defined as:

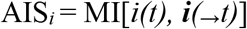

i.e. as shared information between the present activity *i(t)* of a single unit *i* and its past history of activity ***i****(_→_t)* (cf. Figure S6A). Prior to computing mutual information, we binned all spike trains with method as for determining the `Firing`*(t)* descriptive feature vector. The limited amount of available data within each temporal window makes necessary to introduce approximations. Therefore, we replaced the full past history of activity ***i****(_→_t)* with activity at a time in the past *i(t-τ)* and then summed over all the possible lags:

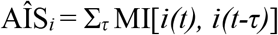

where the lag *τ* was varying in the range 0 ≤ *τ* ≤ 0.5*T_θ_*, where *T_θ_* is the phase of the theta cycle. Note that MI values were generally vanishing for longer latencies (cf. Figure S13A). We evaluated MI terms using a ‘plug-in’ function estimator on binarized spike-trains, which takes the binned spike trains of two neurons for a defined time window and computes the mutual information and entropy values of the two variables (6). Concretely speaking, we estimated the probability *p* that a bin includes a spike and the complementary probability 1 – *p* that a bin is silent for each unit, by direct counting of the frequency of occurrence of “1”s and “0”s in the binned spike trains of each unit. These counts yielded the probability distributions *P(i)* and *P(j)* that two neurons *i* and *j* fire or not. Analogously, we sampled directly from data the histogram *P(i,j*) of joint spike counts for any pair of two units *i* and *j.* These histograms were then directly “plugged in” (hence the name of the used estimator) into the definition of MI itself:

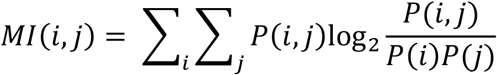

We then subtracted from each MI value a significance threshold (95-th percentile of MI estimated on shuffled binarized trains, 1000 replicas), putting to zero non-significant terms (and thus negative after bias subtraction). Although such corrected plug-in estimator is very rough, it is sufficient in our application in which we are not interested in quantitatively exact values of MI but just in relative comparisons of values, finalized to state clustering over a large amount of observations. We compiled the *N* resulting AÎS*_i_* values into time-dependent vectors `Storage`*(t)*, constructing separate vectors for each of the simultaneously recorded regions. We then constructed storage substates through unsupervised clustering based on the *M_sim_* matrices, as previously described. We defined a neuron to be a *storage hub* in a given state if its AÎS*_i_* value in the state prototype vector was higher than the 95% percentile over all entries of concatenated cluster prototype vectors. Such conservative threshold guarantees that only neurons with exceptionally high AIS values are labeled as hubs. While we may have some false negatives –i.e. neurons with values in the right tail of the AIS distribution not labeled as hubs–, we are thus protected against false positives.

AIS absolute values varied widely between the different recordings. To compare AIS measures and their relative changes between global oscillatory states across recordings, we first averaged AIS for all the units within a specific anatomic layer. We then normalized these average AIS values by dividing them by the average AIS value in the SO state (in anesthesia) or the nonREM state (in natural sleep) for the specifically considered recording and layer. The results of this analysis are shown in Figure S7, where different lines correspond to different recordings.

### Information sharing networks and strengths

Within each time-window we computed time-lagged Mutual Information MI[*i(t), j(t-τ)*] between all pairs of spike density time-series for different single units *i* and *j* (evaluated via the same binning method for determining the `Firing`*(t)* descriptive feature vector). Although MI is not a directed measure, a pseudo-direction of sharing is introduced by the positive time-lag, supposing that information cannot be causally shared from the future. Thus, for every directed pair of single units *i* and *j* (including auto-interactions, with *i* = *j*), we defined pseudo-directed information sharing as:

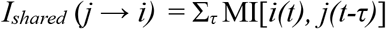

where the lag *τ* was varying in the range 0 ≤ *τ* ≤ 0.5*T_θ_*, where *T_θ_* is the phase of the theta cycle. Once again, we estimated MI terms via direct plug-in estimators on binarized spike trains, as with storage, subtracting a significance threshold (95-th percentile of MI estimated on shuffled binarized trains, 400 replicas) and zeroing not significant terms. All these *I_shared_* (*j* → *i)* entries were interpreted as weights in the adjacency matrix of an information sharing directed functional network, and we defined as *sharing assembly* formed by a neuron *i* the star-subgraph of the information sharing network composed of *i* and all its immediate neighbors. We compiled all the overall *N*^2^ different values of *I_shared_* (*j* → *i)* into time-dependent feature vectors `Sharing_A`*(t)*, describing thus all the possible sharing assemblies at a given time. We then also computed *information sharing strengths* by integrating the total amounts of information that each single unit was sharing with the past activity of other units in the network (“sharing-in”):

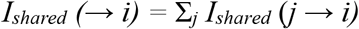

or with the future activity of other units in the network (“sharing-out”):

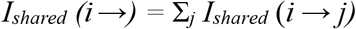

In other words, the integrated amount of shared information was given by the in-strength and the out-strength of a node in the information sharing network with individual link weights *I_shared_* (*j* → *i)*. We compiled the *N* incoming *I_shared_ (*→ *i)* and *N* outgoing *I_shared_ (i* →) values into time-dependent vectors `Sharing_S`*(t)*. We computed separate `Sharing_A`*(t)* and `Sharing_S`*(t)* for each of the simultaneously recorded regions. We then performed as before unsupervised clustering based on the associated *M_sim_* matrices to extract sharing substates. Since the block structure displayed by thee *M_sim_* matrices for sharing assemblies and strengths are nearly perfectly overlapping we conducted all substate analyses based on `Sharing_S`*(t)* vectors only. We defined a neuron to be a *sharing hub* in a given state if its *I_shared_ (_*_* → *i)* and/or *I_shared_ (i* → *_*_)* values in the state prototype vector were higher than the 95% percentile of all concatenated cluster prototypes entries (again protecting against false positive detection).

The relative comparisons of information sharing between SO and THE (REM and nonREM) epochs for different recordings shown in Figure S9, are based, as in the case of AIS in Figure S7, on averaged and scaled values. We first averaged the total *I_shared_* (i.e. sharing in plus sharing out) over all the units within a specific anatomic layer. We then normalized these average total *I_shared_* values by dividing them by the average total *I_shared_* value in the SO state (in anesthesia) or the nonREM state (in natural sleep) for the specifically considered recording and layer.

### Liquidity of sharing

The *M_rec_* matrices for Sharing Assemblies display light blue (low internal correlation) blocks while the *M_rec_* matrices for Sharing Strengths have similar blocks but red-hued (higher internal correlation). We quantify this visual impression by evaluating liquidity of sharing strength and sharing assembly substates. For a given recording and a given associated *M_rec_* matrix (e.g. the one for the `Sharing_A` or the `Sharing_S` features), we define *T*_α_ as the set of times *t* for which the system is in a given substate α relative to the considered feature of interest. We then evaluate the liquidity Λ(*α*) of this substate α as:

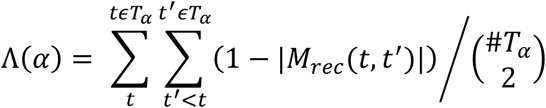

where | · | denotes the absolute value operator and #*T*_α_ is the number of elements of the set *T*_α_. Liquidity values are thus bounded in the interval 0 ≤ Λ(*α*) ≤ 1, with 1 indicating maximum liquidity (i.e. maximum internal variability) of a substate.

### Oscillatory mode specificity and hub distributions

For each substate (firing, storage, sharing) we computed the fraction of times that the substate was observed during a SO or a THE state (in anesthesia) or a nonREM or REM state (in natural sleep). We defined the largest among these fractions as the oscillatory specificity of this substate. Oscillatory specificities close to 1 indicate that a substate occurs mostly within one of the two possible global states observed in each recording, while specificities close to 0.5 indicate that the substate do not occur preferentially in one of the global states.

To evaluate the probability that a hub emerges in a given anatomical layer, we computed for every recording the fraction of cells recorded in each layer that were labeled as hubs at least in one computing substate (storage or sharing). We computed separately these fractions layer-by-layer, for excitatory and inhibitory cells and for anesthesia or sleep. These fractions were equal to unit when all the excitatory (or inhibitory) cells in a layer happened to be hubs at least once. We then evaluated the general probability that a hub emerges in a layer, which is different from the previous one, because it takes in account as well the fact that some cells may be labeled as hubs more often than others. We then considered the list of all hubs of a given type (storage or sharing) across all substates, including repetitions (if a neuron was hub in more than one substate then it appeared multiple times in the list) and evaluated the fraction of times in which a hub in this list was belonging to a given layer. We computed separately these fractions layer-by-layer, for storage or sharing hubs and for anesthesia or sleep. 95% confidence intervals for the mean fractions above were evaluated as 2.996 times their sample standard deviation over the different recordings for which they could be computed. We considered two mean fractions to be different when their 95% confidence intervals were fully disjoint.

### Coordination between substate transitions

To compare sequences of substates of different types or in different regions we introduced a symbolic description of substate switching. Each substate was assigned a *letter* symbol, i.e. a label *s^(p)^* where *p* can stand for firing, information storage or sharing and *s^(p)^* is an arbitrary integer label different for every substate. We could thus describe the temporal sequences of the visited substates of each different type as an ordered list of integers *s^(p)^(t)*. We quantified the degree of coordination between the sequences of substates of different types (e.g. *p* = ‘storage’ vs *q* = ‘sharing’) or in different regions (e.g. *p* = ‘storage in EC’ vs *q* = ‘storage in CA1’) by evaluating the relative Mutual Information term:

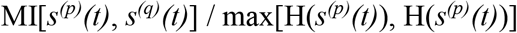

normalized between 0 –full statistical independence between the two substate sequences– and 1 –complete overlap between the two substate sequences–, by dividing it by the entropy H of the most entropic among the two symbolic streams. We evaluated these MI and H terms using direct plug-in estimators on the joint histograms of substate labels. We estimated chance expectations for the level of coordination by repeating the same procedure for substate sequences with shuffled substate labels and then finding the 99^th^ percentile over 1000 permutation replicas of the computed MI/H.

### Mutual Information Measure’s Dependence on Bin Size

The original decision for the bin size was chosen such that when discretized, the information content lost by counting 2 or 3 spikes on the same neuron within a given bin as a ‘1’ was less than 5%. On average, the information content lost was less than 1% across all recordings. To analyze the dependence on bin sizes, one example recording was chosen in the PFC during natural sleep in different bin sizes, 25 *ms*, 33 *ms* and 66 *ms* and computed substates using the same methods described above. To make the comparison focused on bin size, the same number of clusters per feature was chosen to reflect the original number. We then computed the amount of information about the substate sequences computed with the original binsize were retained by corresponding substate sequences derived for each different bin size. To do so, we used the same procedure described in the previous section to quantify coordination between sequences for different types of states or between different regions. Notably we computed mutual information between the substate sequences for different bin-sizes (normalized by the entropy of the original sequence) and compared this relative mutual information with chance expectation (obtained via shuffling substate sequences, as above). We found that the mutual information between corresponding sequences for different bin sizes was two order of magnitudes above chance level (Figure S13B), denoting high robustness of our procedure for extracting substates. Correspondingly, we also found that, for matched substates between sequences extracted for different bin-sizes, the identification, number and anatomical localization of hubs were only marginally altered.

### Complexity of substate sequences

After converting sequences of substates into symbolic streams of letters, we defined substate *words* as the triplets of letters corresponding to the firing, the information storage and the information sharing substates simultaneously observed at each time *t*, i.e.:

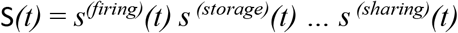

We then constructed a *switching table* T in which the temporally ordered columns provide the sequence of substate words S*(t)* along time. We compiled separate switching tables for each recording and for each of the simultaneously recorded regions. The total set of substate words effectively found in a specific switching table constitutes its associated *dictionary* of substate combinations. We defined then the *used dictionary fraction*, as the ratio between the number of observed words and the maximum theoretically possible number of words that could have been composed given the available substate letters (depending on how many firing, storage or sharing substates have been effectively extracted).

We then evaluated the complexity of substate word sequences using a procedure inspired from the notion of Kolmogorov-Chaitin complexity (*19*) and minimum description length approaches (MDL; *20*). The basic concept is that, for a regular symbolic sequence (as our streams of substate words), it will be possible to design a tailored “compression language” such the sequence will admit a much shorter description when reformulated into this language with respect to the original length in terms of number of words. On the contrary, a random symbolic sequence will be poorly compressible, i.e. its descriptions in terms of a generative language will be nearly as long as the original list of symbols appearing in the sequence. A complex symbolic sequence will stand between these two extremes - still admitting a compressed generative description but not as short as for regular sequences. Departing from universal compression approaches, as the original MDL formulation (*20*) or the Lempel-Ziv approach (*21*), we introduce here a “toy language” for generative description, specialized to compress state transition tables as the ones of Figure 6A. Our choice is conceptually compliant with the MDL approach but – for the sake of pedagogy– avoids technical steps as the use of binary prefix coding.

Let Ω ={S_1_, S_2_,…, S_ω_} be the dictionary of substate words appearing in the switching table T which we want to describe. We first define the *exhaustive list description* (*D*_list_) of T as a string of the following form:

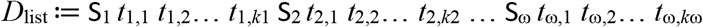

In such a description the symbol of each substate word S*_q_* (counting as one description unit) is followed by an exhaustive list of all the *k_q_* times *t_q_*_,1_, *t_q_*_,2_,…, *t_q_*_,*kq*_ (each time index counting as an extra description unit) at which the recorded system produced the matching substate word. If the number of analyzed time windows is *K* = *k*_1_ + *k*_2_ +…+ *k*_ω_, then the length of the exhaustive list description will be |*D*_list_| = *K* + ω description units (*K* time stamps, plus ω substate word symbols).

Let then define the *block-length description* (*D*_block_) of the stream of substate codewords, as a description of the following form:

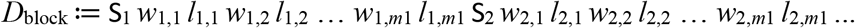

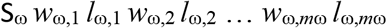

In such a description the symbol of each word S*_q_* (always counting as one description unit) is followed by a list of stepping instructions for a hypothetical “writing head” moving along different discrete positions on an idealized tape, similarly to computing automata as the Turing Machine (*69*). At the beginning the machine is initialized with the head on the first position on the tape. The integers *w_q_*_,*n*_ –at odd positions (1^st^, 3^rd^, etc.) after the word symbol– indicate for how many steps the machine head must shift on the tape toward the right *without* writing, but just skipping positions. The integers *l_q_*_,*n*_ –at even positions (2^nd^, 4^th^, etc.) after the word symbol– indicate instead for how many steps the machine must also write on the tape the symbolic string S*_q_* before then shifting to the next position on the right. Every time that a new symbol S*_q_* is met when parsing the step lengths description, the position of the writing head is reset to the leftmost starting position on the tape. Such parsing grammar is obviously more complex than the one for a simpler “parrot machine”, designed to parse exhaustive list descriptions as the ones described above. The length in symbols of this block-length description is variable and depends on how regular the word sequence is to compress and regenerate. The block-length description segment S*_q_ w_q_*_,1_ *l_q_*_,1_… *w_q_*_,*mq*_ *l_q_*_,*mq*_ will be shorter than the matching exhaustive list description segment S*_q_ t_q_*_,1_… *t_q_*_,*kq*_ whenever 2*m_p_* < *k_p_*, which can happen if transitions for the different types of substate letters are regularly aligned, in such a way that the resulting switching table have long alternating blocks with repeated substate words.

The syntactic complexity of a sequence of substate words can then be evaluated by quantifying how much the program to generate the switching table T via a “smart” compressing machine interpreting block-length descriptions is shorter than the program to generate the same table T via a “dumb” parrot machine interpreting exhaustive length descriptions. We define the *description length complexity* of a switching table T as:

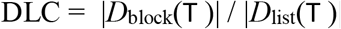

To give a toy example, let’s consider the sequence T = “AAAAAAA BBBB AAAAA CCCCC DDD BBBBBB”, built out of four possible collection of substate words S_1_ = A, S_2_ = B, S_3_ = C and S_4_ = D. The exhaustive list description for this sequence will be:

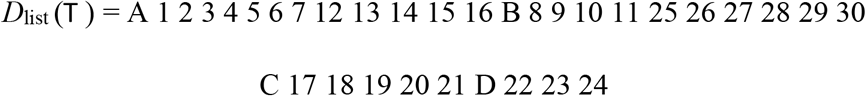

with length |*D*_list_(T)| = 34 descriptive units. Its step lengths description will be:

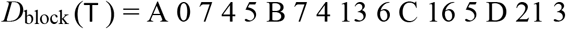

with length |*D*_block_(T)| = 16 descriptive units, i.e. |*D*_block_(T)| < |*D*_list_(T)|.

Given the noisiness of data, we dropped from both the exhaustive list description and the step lengths description the segments corresponding to exceedingly rare words S*_q_*. In particular, ranking the code words from the least to the rarest, we dropped all the words S*_r_* with *r* ≥ *R*, such that removing all of their occurrences in the word stream reduced the stream’s overall length of no more than 10% (lossy compression).

We computed confidence intervals for DLC values via a Jacknife construction in which we drop one word at random position from the temporal stream S*(t)* made of *K* symbols, generating up to *K* Jackknife surrogate streams, each with *K*-1 symbols. The confidence interval was then given by the 5-th and the 95-th percentile over the complexities evaluated from these Jackknife surrogates.

Appropriate reference criteria were then required to discriminate complex vs ordered or random switching tables. We need to compare the empirically observed DLC values against two thresholds. Complex switching tables should have indeed a DLC below a threshold for randomness testing and above a threshold for regularity testing. Given a switching table T, we constructed a randomized version T_rand_ by randomly permuting independently the entries of each of its rows. For each recording, we constructed 1000 instances of T_rand_ and evaluated DLC for all of them, identifying as upper threshold for complexity the 5-th percentile DLC_rand_ = *q*_5%_[DLC(T_rand_)]. We then constructed an enhanced-regularity version T_regular_ of each T by lexicographically sorting entries row-by-row (to get blocks of homogeneous code-words as long-lasting as possible based on exactly the same building bricks). We then arbitrarily defined a lower threshold for complexity DLC_regular_ = 2 DLC(T_regular_). The thresholds DLC_regular_ and DLC_rand_ varied for every switching table. However, the criterion DLC_regular_ < DLC < DLC_rand_ was fulfilled for all the considered recordings, whose state transitions sequences could then be certified to be complex (in our arbitrary but quantitative and operational sense).

Importantly, we could restrict the evaluation of complexity to sub-table restricted to words occurring during selected different global oscillatory states only. In this way we could compare the complexity of sequences occurring within different global states, e.g. REM vs nonREM. We plot in Figure 6E relative complexity variations between two global states α and β:

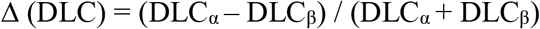

We evaluated once again confidence intervals for relative complexity variations via one-leave-out Jacknife on global state-restricted switching table columns.

### Burstiness of state sequences

We also characterize switching tables in terms of their “style” of transitions, looking at two different temporal statistics. First, we computed all *inter-transition times* from a table T, i.e. the number of time-steps occurring between one block (continuous time-interval with a same substate word maintained in time) to the next. Note that these inter-transition times are precisely the *l_p_*_,*n*_ integers appearing in the block-length description *D*_block_ (T) of the table T. After computing the mean *μ*_*l*_ and the standard deviation *σ*_*l*_ of these inter-transition times, we then evaluated the burstiness coefficient (*22*):

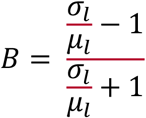

Such a coefficient is bound between −1 < B < 1 and is equal to 0 when transitions between substate words follow a Poisson statistic, negative when the train of transitions is more periodic and positive when more bursty than for a Poisson train.

## Supplementary figures

**Figure S1.**
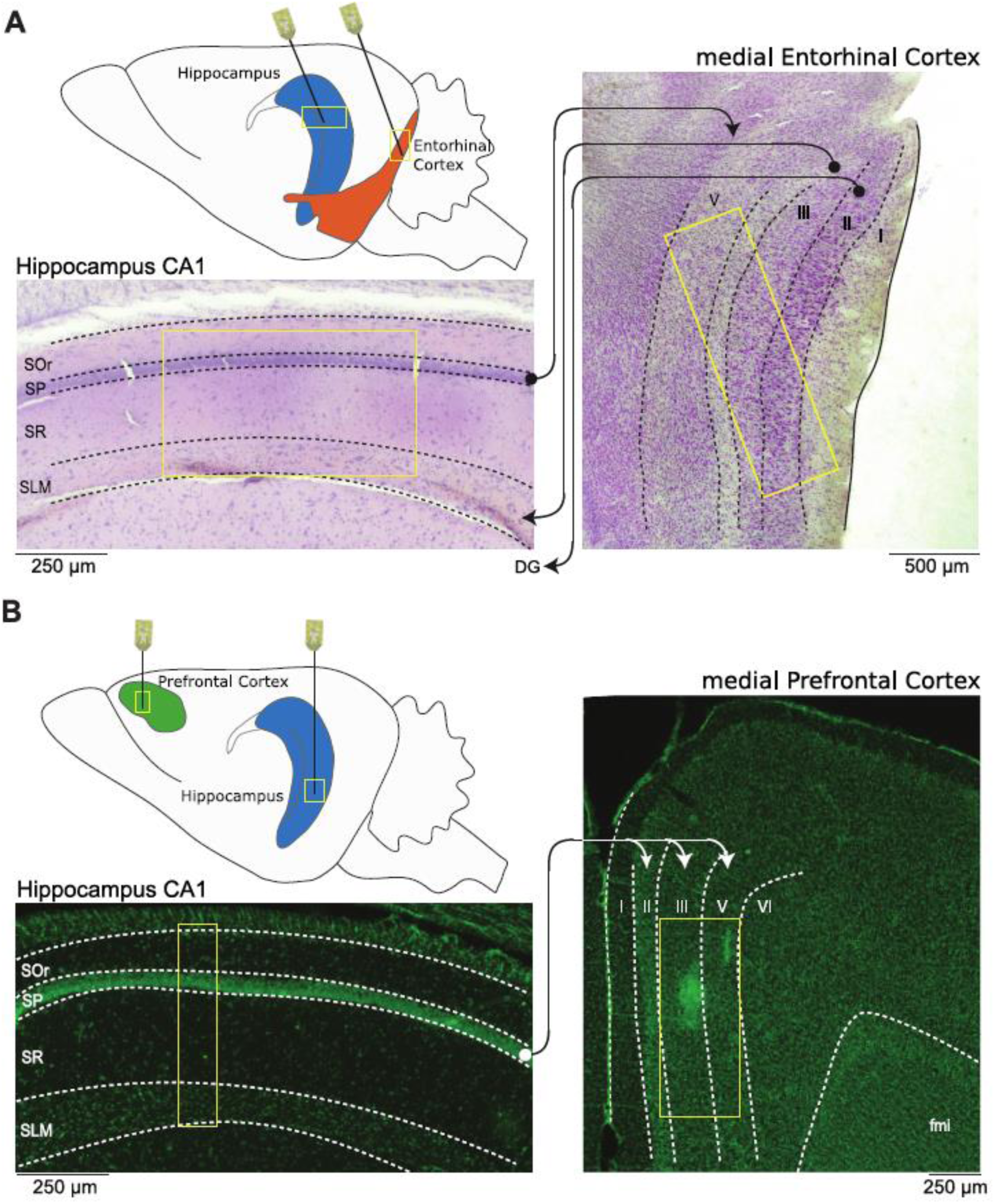
Recording paradigm. Schematic representation of the (**A**) simultaneous mEC/HPC recording setup and (**B**) simultaneous mPFC/HPC during anesthesia and natural sleep. The Nissl stained sections display the anatomical regions recorded by the different silicon probes used (yellow boxes). Arrows represent the anatomical connectivity (●: source layer, ➔: target layer) between the dorsal hippocampus CA1 region (SOr: stratum oriens; SP: stratum pyramidale; SR: stratum radiatum; SLM: stratum lacunosum moleculare) with the dorso-medial entorhinal cortex (mEC, layers I to VI) and medial prefrontal cortex (mPFC, layers I to VI).

**Figure S2.**
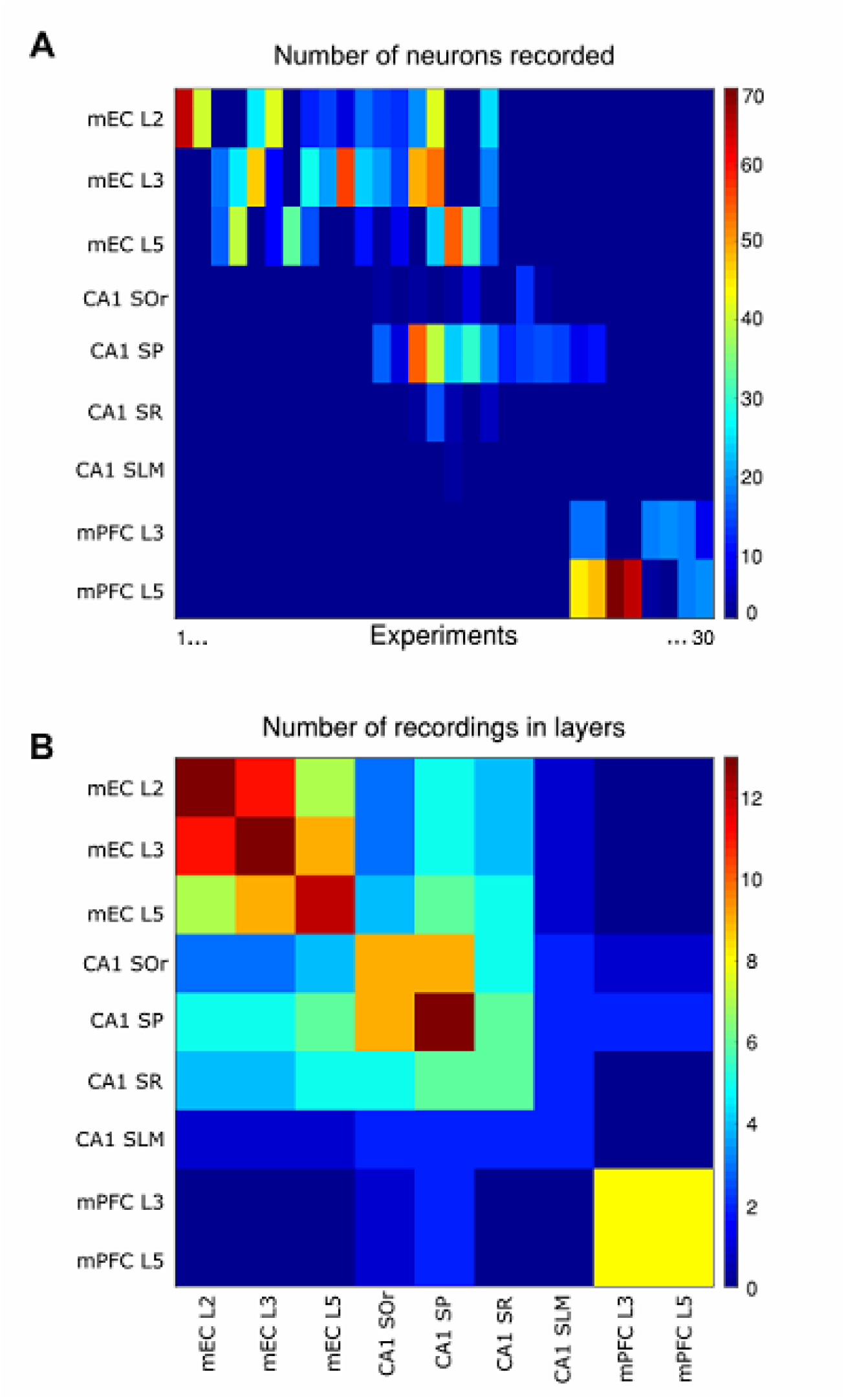
Information about recordings. We analyzed data from 30 different recordings performed in 24 different rats. In each recording we identify single units in different anatomical locations. **(A)** Number of recorded single units (color coded on the right scalebar) per anatomical layer (rows), for each of the 30 recordings (columns). **(B)** Number of recordings (color coded on the right scalebar) simultaneously targeting pairs of two different anatomical layers.

**Figure S3.**
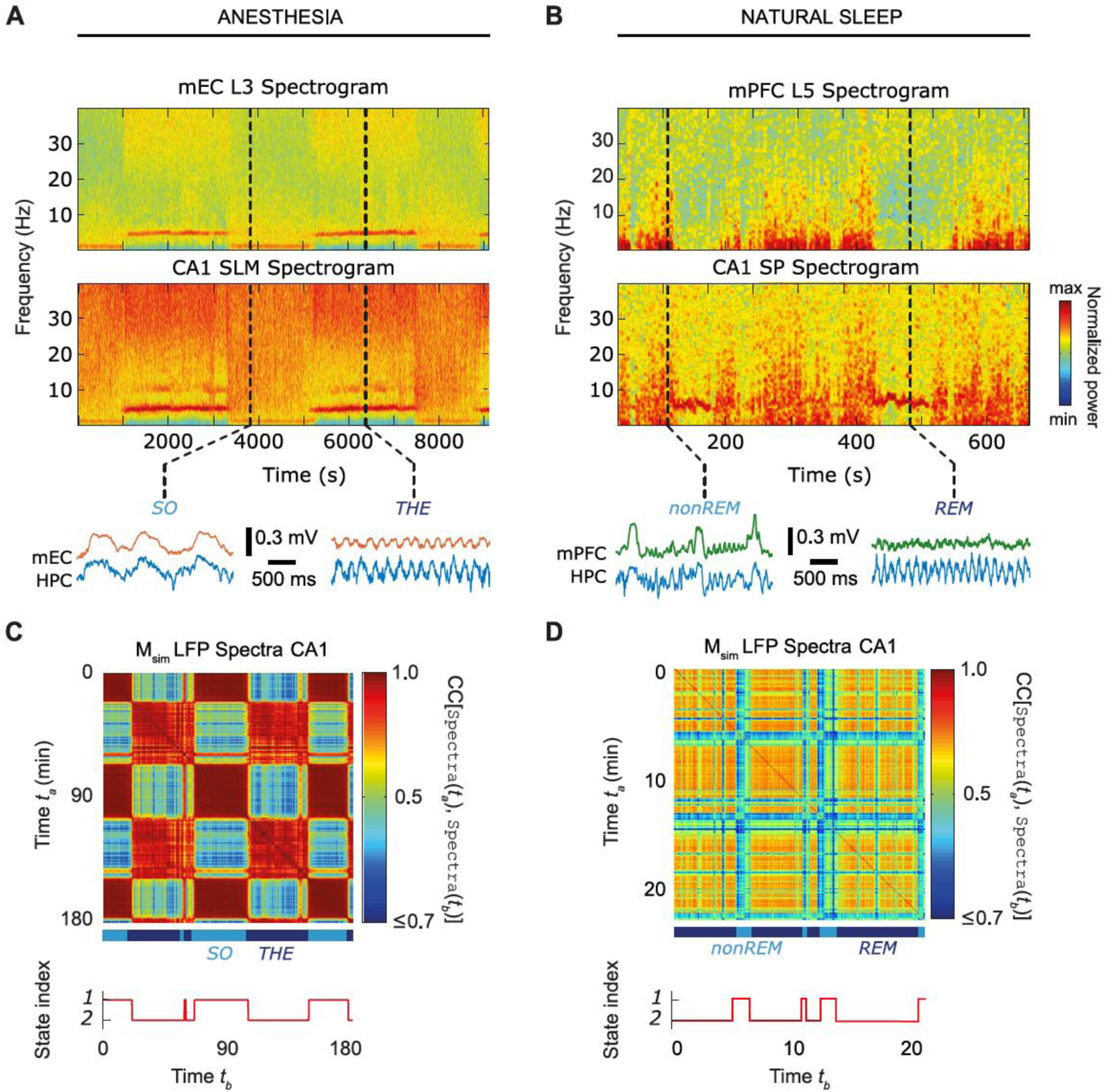
Global brain oscillatory states. We performed a time-frequency spectral analysis of the LFP signals from all channels. Time-frequency spectrograms of LFPs are shown in **(A)** from mEC III and CA1 SLM layers during anesthesia and in **(B)** for mPFC layer V and CA1 SP. A characteristic alternation is visible between epochs dominated by SO/THE rhythms and REM/nonREM. Example LFP traces at time points corresponding to the dashed vertical lines are magnified and shown below the spectrograms. To characterize global oscillatory states in an unsupervised manner within each time-window of analysis (Figure 1), we averaged the power across different frequency bands and compiled all LFP channels into the feature vector `Spectra`*(t)*. The similarity matrices corresponding to (A) and (B) are shown in (C) and (D), respectively. The alternation between SO and THE epochs in **(C)** and between nonREM and REM epochs in **(D)** is well visible in the marked block structure of the feature similarity matrices. Unsupervised clustering identified 2 states under anesthesia or natural sleep.

**Figure S4.**
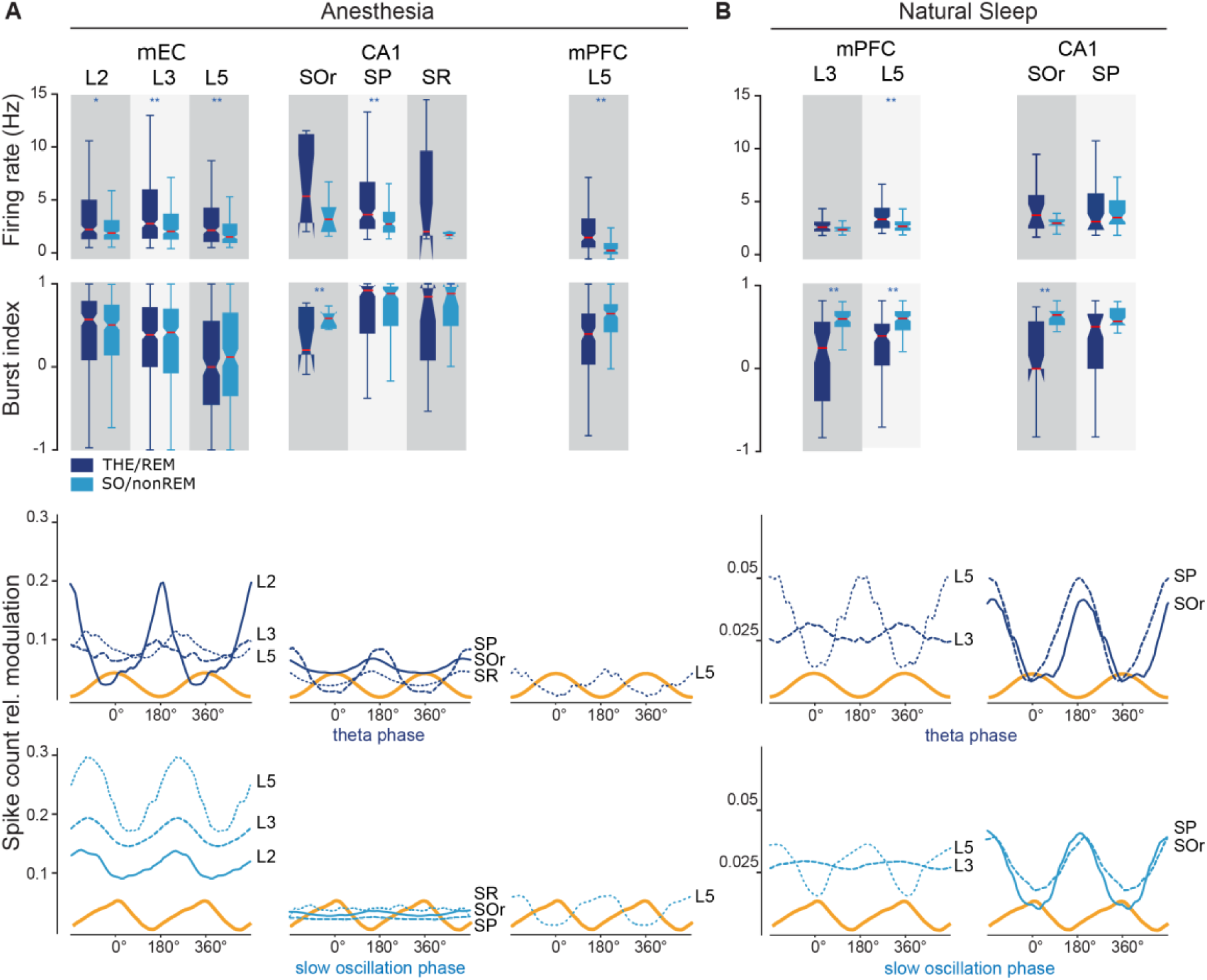
Effects of global states on unit firing. Transitions between global oscillatory states in both anesthesia and sleep significantly modulated the median firing rate in mEC, mPFC and CA1 layers (**A**, top). Burstiness (**B**, bottom) was significantly modulated during natural sleep only. Both firing rate and bursting indices were heterogeneous across neurons, as emphasized by long box-plot whiskers. Single units in mEC, mPFC and CA1 layers fired preferentially at well-defined phases of the ongoing theta (**B**, top) and slow oscillation (**B**, bottom) rhythms as visualized by phase-binned histograms of spike count relative modulation, compared with reference average LFP waveform cycles. Note that phase modulations were an order of magnitude stronger during anesthesia than natural sleep.

**Figure S5.**
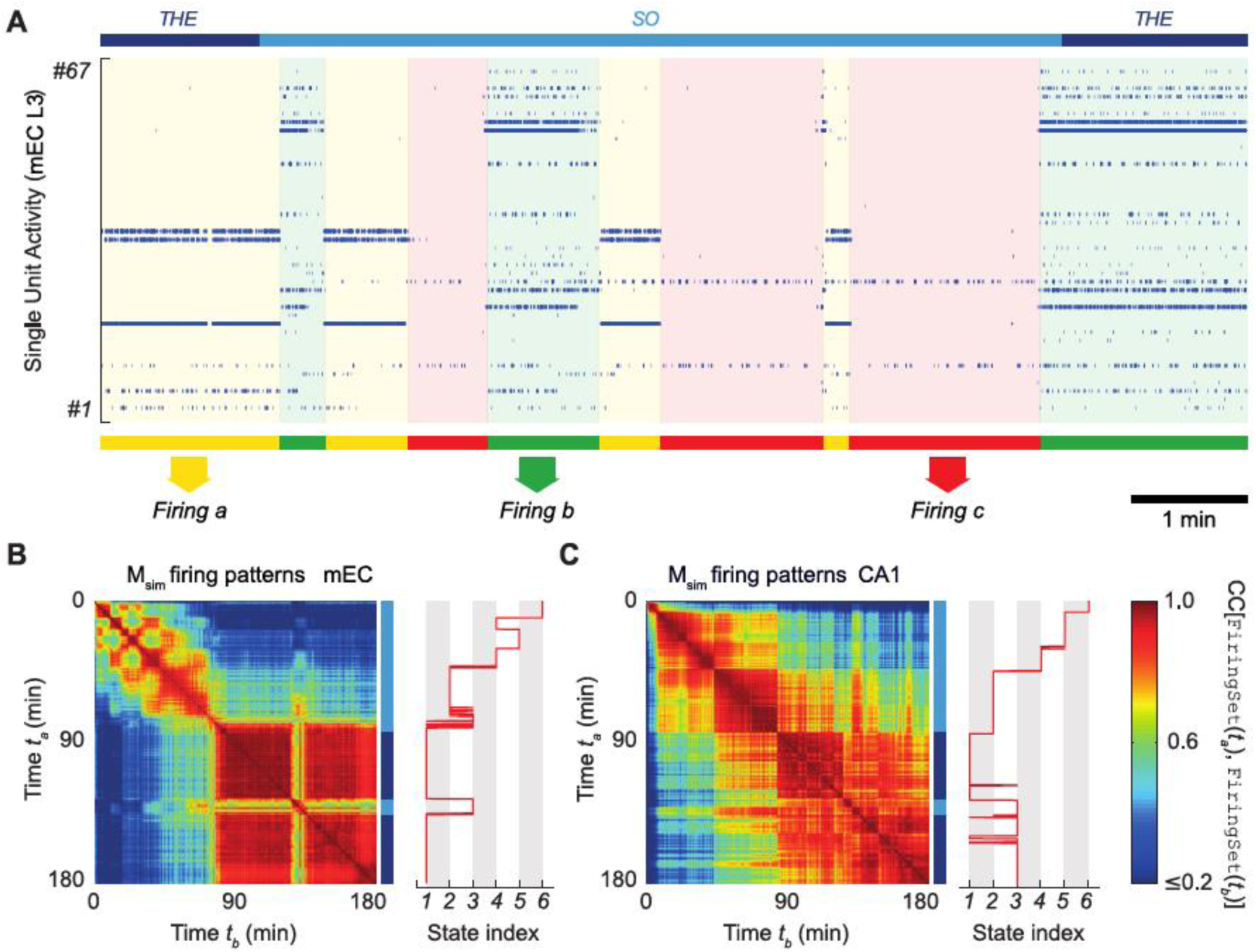
Firing substates. (**A**) displays the firing, represented by blue dots, of 67 neurons recorded in layer 3 in mEC. The solid lines above indicate the global brain states (THE and SO) identified using unsupervised clustering of the spectral features of the field potential (as described in Figure S3). Unsupervised clustering identified three sets of co-firing neurons (indicated by yellow, green and red solid lines), which are clearly visible, during the recording time shown here. Note the alternation of the firing substates, and the fact that global oscillatory state transitions (THE→SO→THE) do not correspond to transitions between firing substates. Below, (**B**) and (**C**) represent the dynamics of firing sets in mEC and CA1 during the whole recording session, where squares across the diagonal represent different firing substates, as in (**A**).

**Figure S6.**
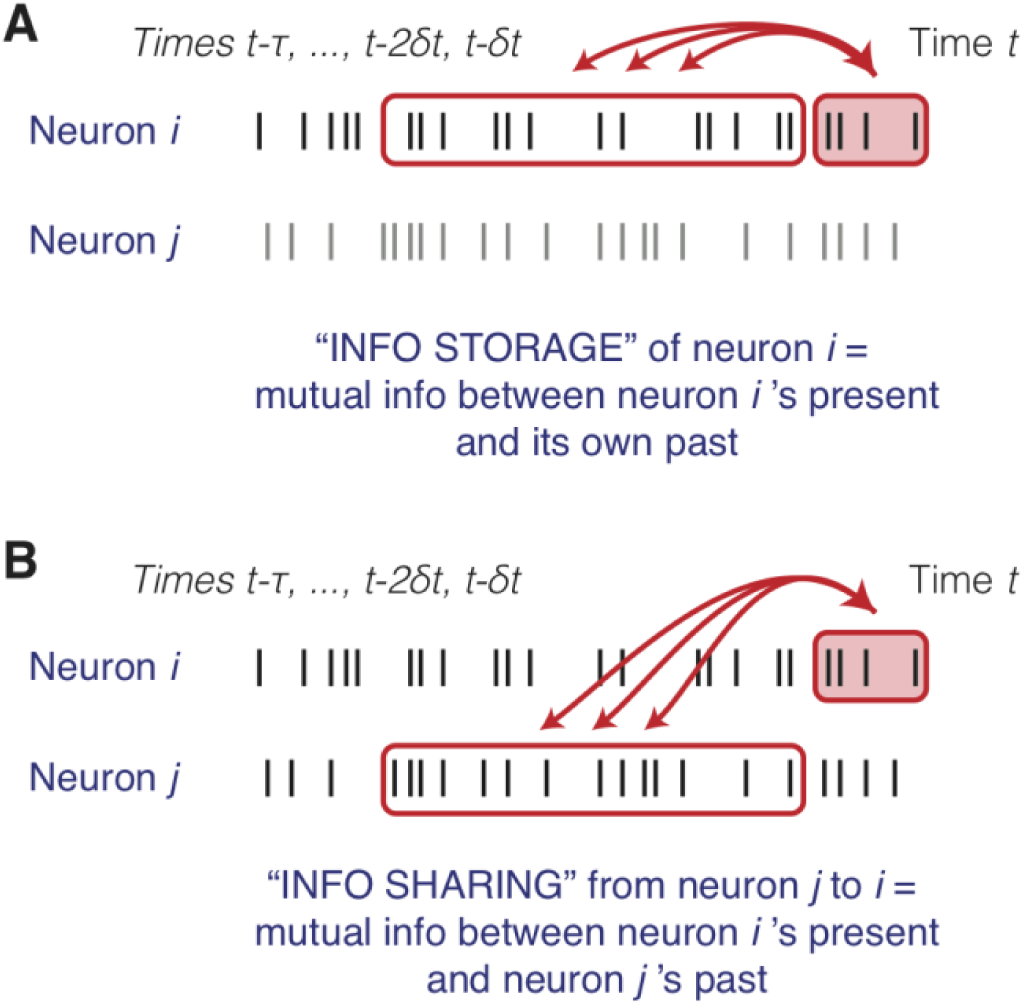
Primitive operations in information processing: explanatory cartoons. The information conveyed by the activity of a unit at a given time may have different sources. A fraction of the total information conveyed at time *t* by a neuron *i* may have already been present in *i*’s past activity (**A**). Therefore, we say that this fraction of information is being *actively stored* into neuron *i* by its activity, implementing a “memory buffer”. The involvement of a unit into this primitive information processing operation is quantified by its Active Information Storage score (see Figure 3). A complementary fraction of the total information conveyed at time *t* by a neuron *i* may have been present already into the past activity of a different neuron *j* (**B**). We say in this case that this fraction of information is *shared* from *j* toward *i*, with time-lagged mutual information providing a pseudo-directed measure of functional connectivity (see Figure 4).

**Figure S7.**
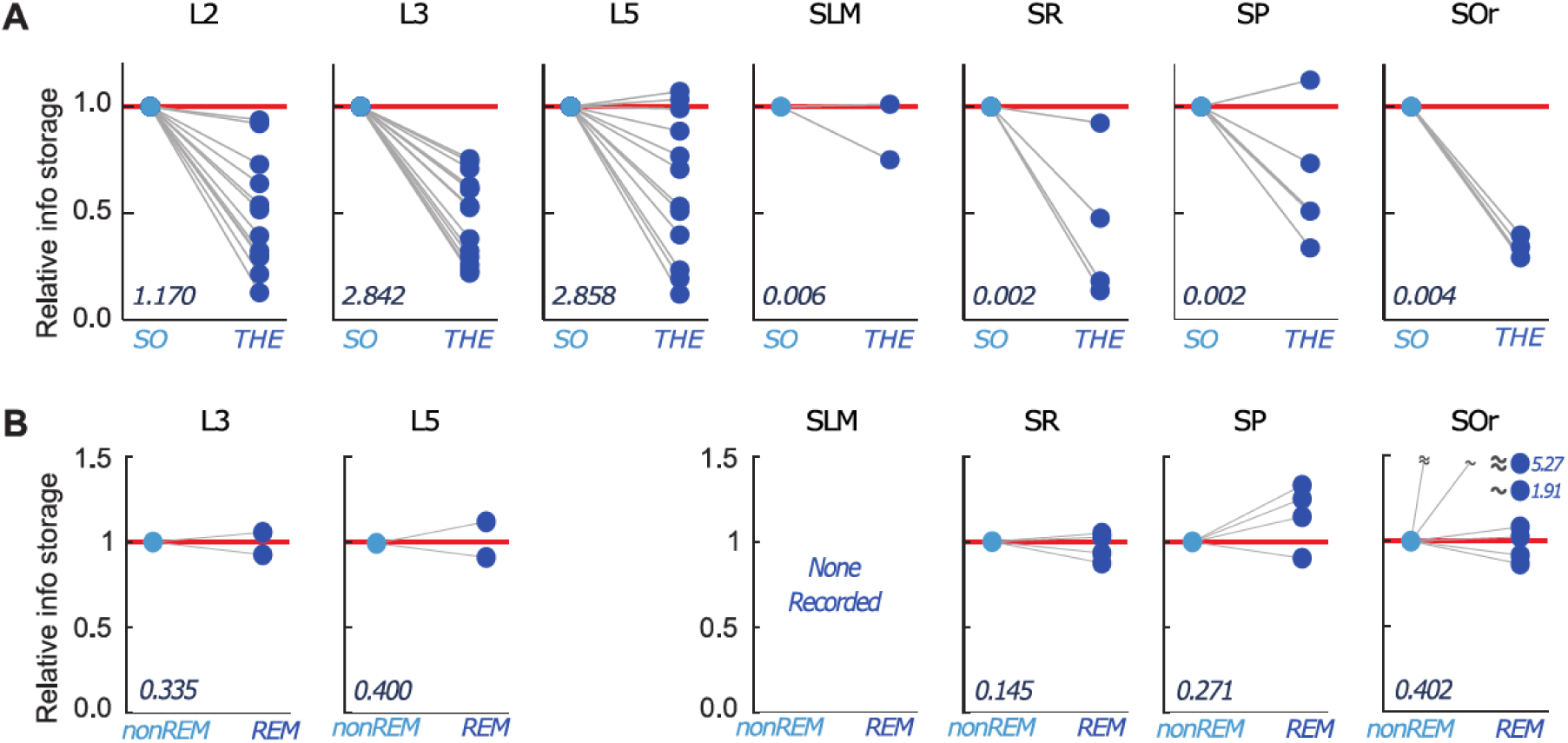
Information storage is brain state-dependent. **(A)** Relative variation of active information storage values in the different mEC and CA1 layers between SO and THE states during anesthesia. Different lines correspond to different rats. All values are normalized to SO values for better visualization of the size and direction of effects. The absolute values of active information storage during SO are indicated in the lower left corner of each subpanel. During anesthesia, mEC layers have larger active information storage values than CA1 layers. Generally, switching from SO to THE state tends to reduce active information storage values. **(B)** Same as A but for mPFC and CA1 layers during natural sleep. The active information storage has the same order of magnitude for mPFC and CA1 but is lower than mEC during anesthesia. Furthermore, during natural sleep there is no major general difference between REM and nonREM (as opposed to SO/THE in A).

**Figure S8.**
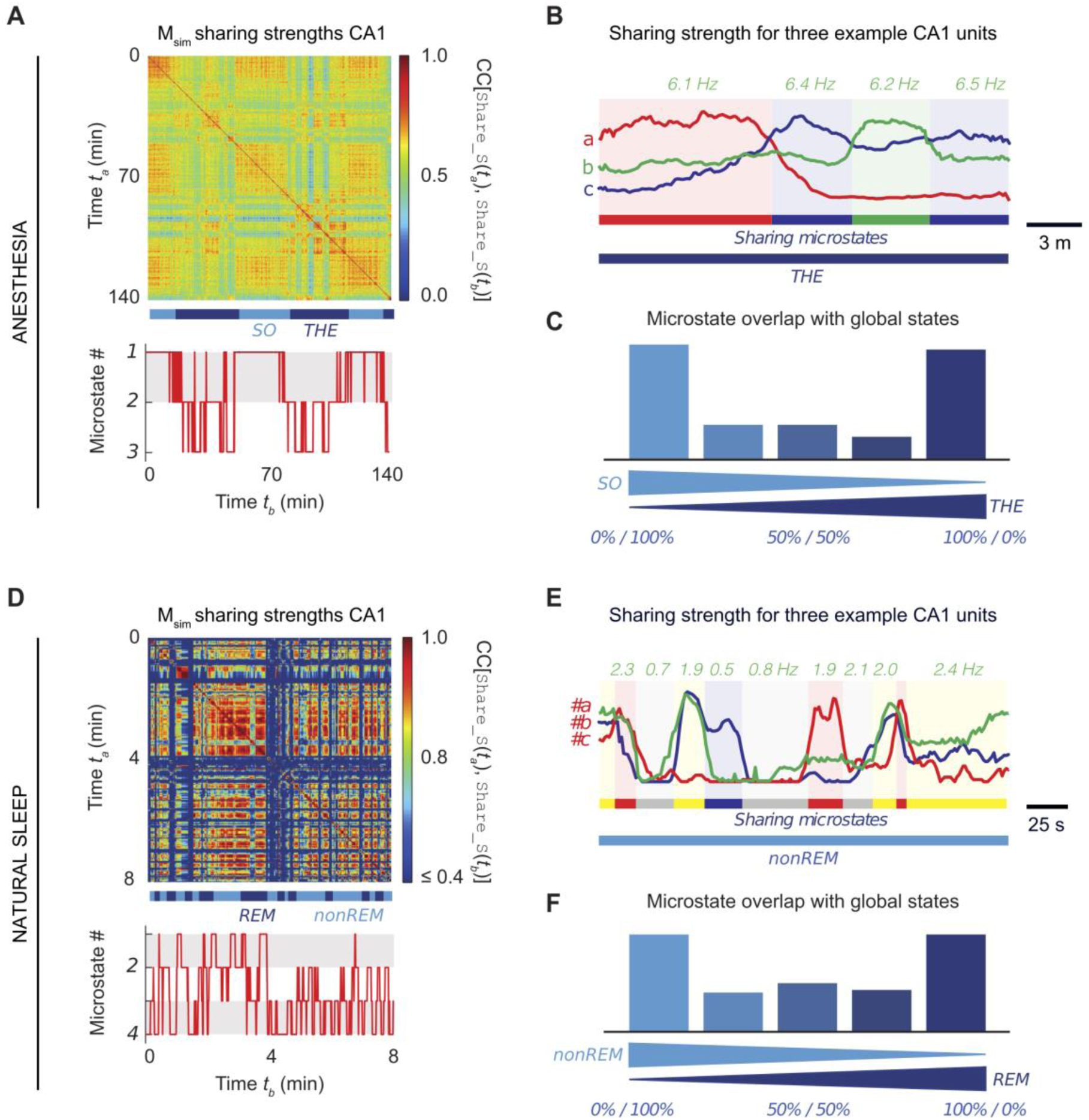
Substates of information sharing: additional information. We show here additional typical feature similarity matrices *M_sim_* obtained from sharing strengths `Sharing_S`*(t)* in CA1 during anesthesia **(A)** and natural sleep **(D)** (see also Figure 4). As in firing and storage substates, **(C)** and **(E)** show that the participation of different neurons to the information sharing was varying along time in a switching fashion (arbitrary normalized units, smoothed time-series). The values reported above the plots correspond to the average firing rate of the neuron *b* (green color) during the corresponding epochs within consistent sharing substates. Sharing substates tended to occur during a preferred global oscillatory substate, as indicated by the bimodal histograms in **(C)** for anesthesia and **(F)** for natural sleep (see also Figure 4D).

**Figure S9.**
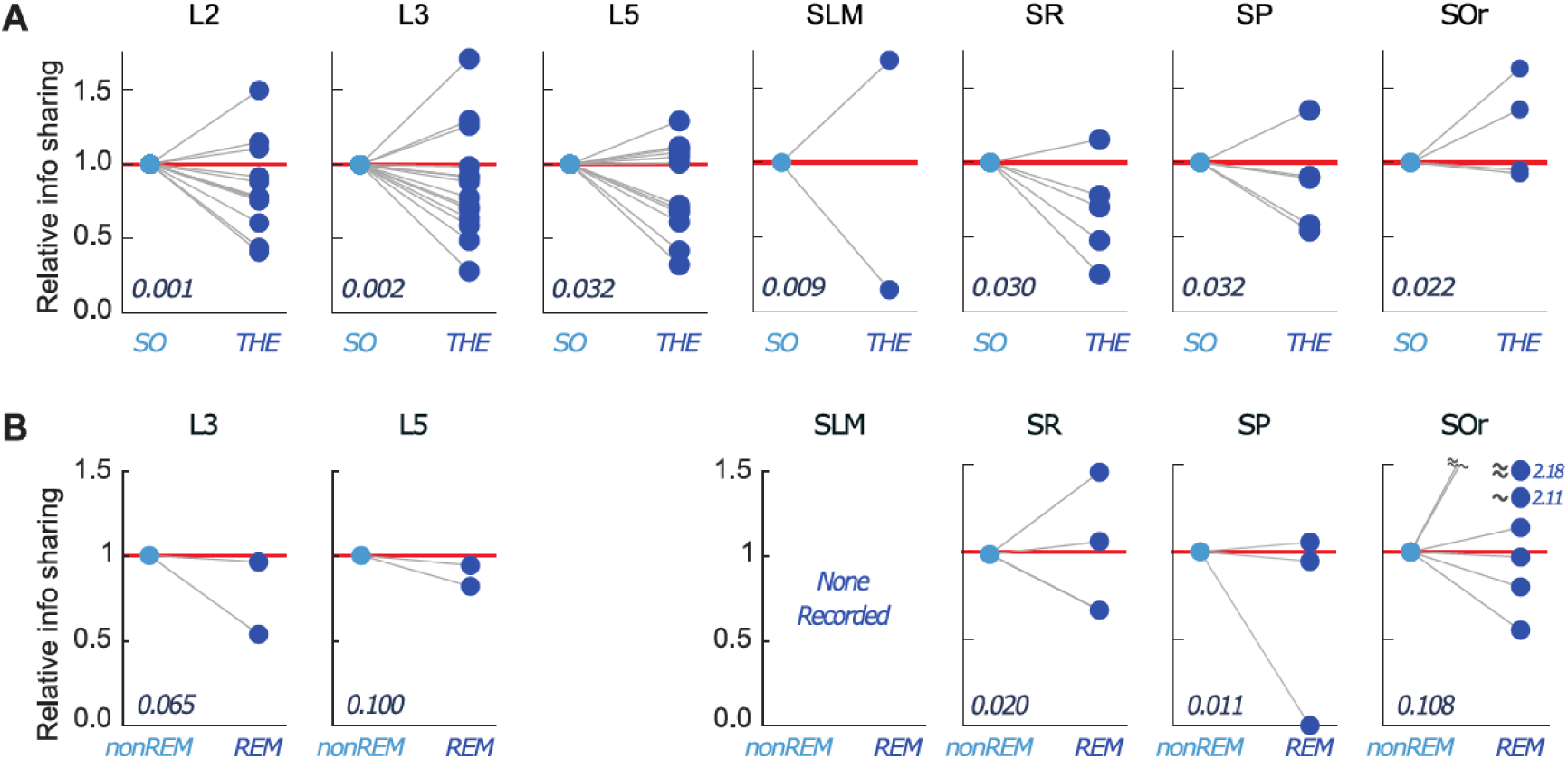
Variations of information sharing as a function of the global brain oscillatory states. **(A)** Relative percent variation of information sharing total strength values between SO/THE states during anesthesia, averaged over different layers in mEC and CA1. Different lines correspond to different rats and average values of sharing strength in SO state are normalized to allow a simpler comparison of the size and direction of effects for different layers, but absolute values of sharing strength in the SO state, averaged over the different rats, are indicated in the lower left corner of each subpanel. **(B)** Same as above but for different mPFC and CA1 layers during natural sleep. We did not observe any systematic direction of change for sharing strengths when switching from SO to THE states during anesthesia or from nonREM to REM during natural sleep. mEC layers II and III during anesthesia were associated to the weaker absolute values of sharing strengths and mPFC layer IV and CA1 SO during natural sleep to the stronger values.

**Figure S10.**
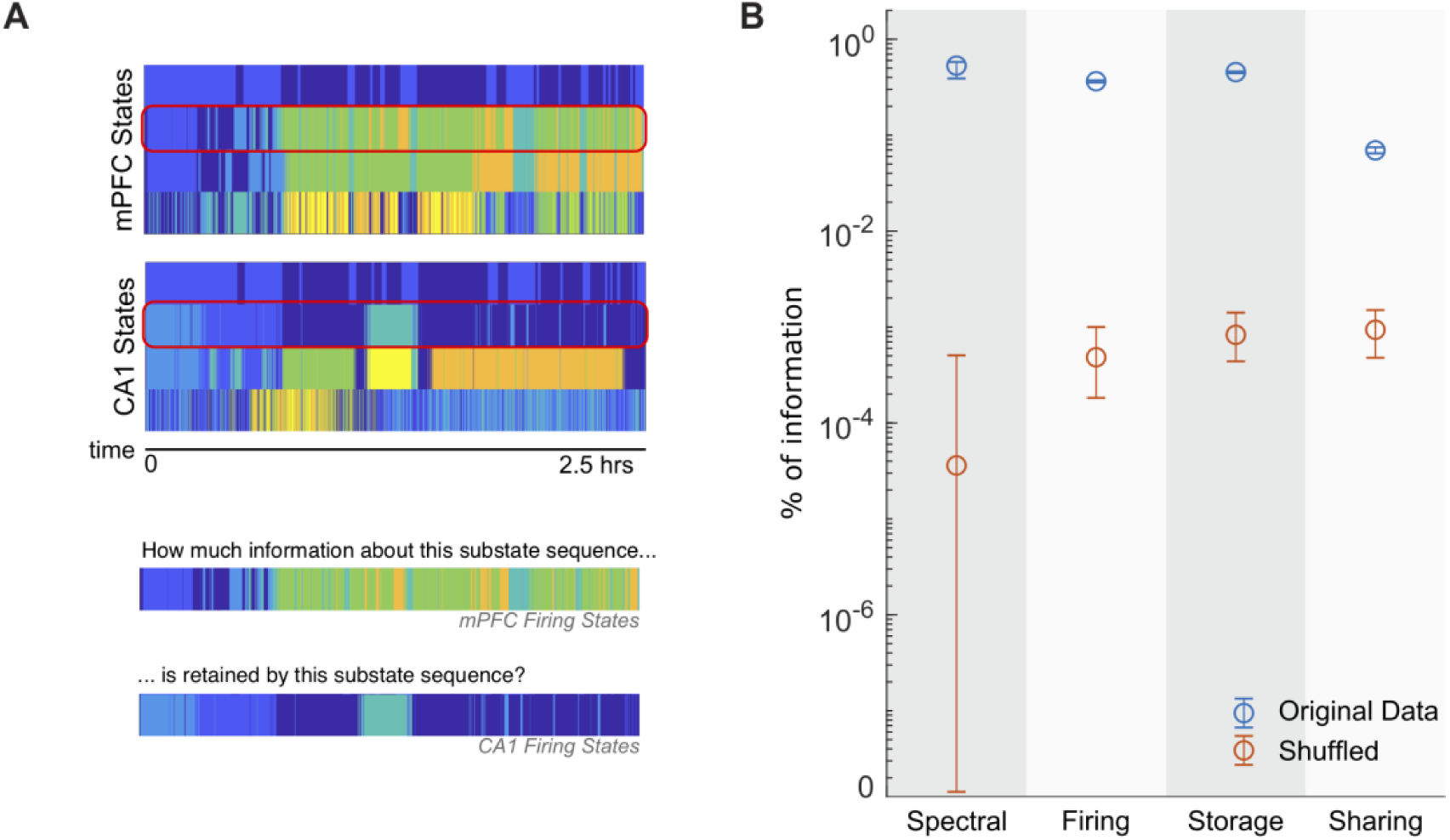
Coordination of substate transitions between brain regions. Sequences of firing, storage and sharing substates do not show a perfect match between simultaneously recorder regions, as visible from the state transition tables of a representative simultaneous CA1/mPFC recording during natural sleep (**A**, top). To quantify the level of interregional coordination between substate transitions we evaluated the mutual information between matching substate sequences in the two simultaneously probed regions (**A**, bottom). We also estimated the corresponding chance levels of coordination, by repeating the same procedure on shuffled state transitions sequences. For all features, we find that the sequences are loosely coupled between regions, but still far above chance level (**B**). Vertical bars denote the 99% confidence interval (bootstrap with replacement for original data, permutation-based for shuffled data, 1000 replicas in both cases). This specific graph is built using the example in (**A**).

**Figure S11.**
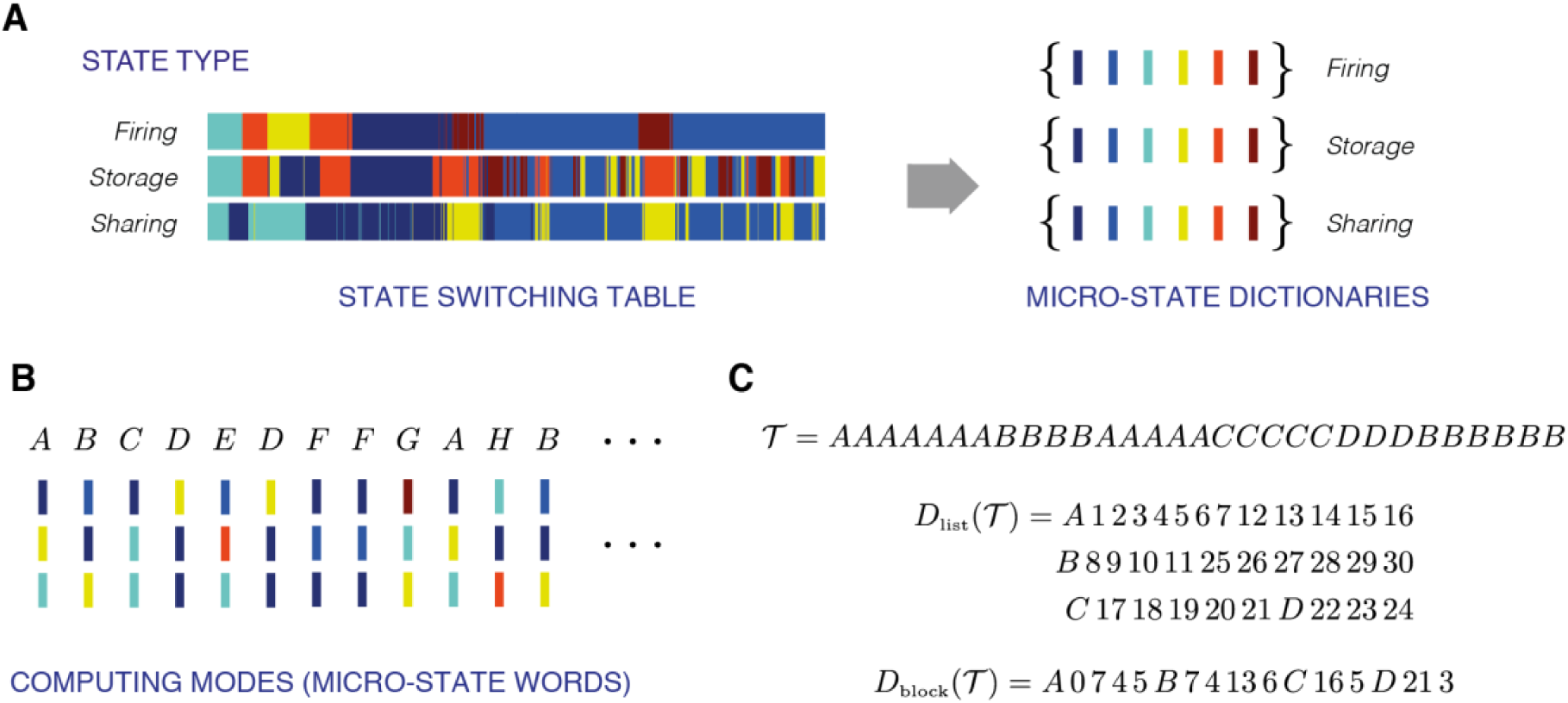
Calculation of complexity of the state switching table. **(A)** A given recording can be represented by the superposition of 3 sequences of substates for each feature (firing, storage, sharing). **(B)** At any point in time, the triplet of substates can be represented as a *word* made of three substate *letters*. For simplicity, the *letters* are color-coded, and a *word* is represented by a letter from the alphabet (A, B, C etc.). **(C)** A state switching table *𝒯* is the temporal sequence of *words* with one word per analysis window *t_A_* as defined in Figure 1). There are two ways to represent the sequence of *words*. *D*_list_*(𝒯)* lists the positions at which the *words* appear (e.g. at window 1, 2, 3, 4, 5, 6, 7, 12, 13, 14, 15, 16…). This representation is exhaustive, but not compact. *D*_block_*(𝒯)* lists how many positions one should skip from the start of the sequence before writing and in how many consecutive positions the considered *word* should be printed. In the example shown here, the *word* “A” occurs at the beginning of the string – zero positions skipped – and is then printed seven times. After, 4 positions are skipped, and it is written again 5 times, etc. This representation is more compact. Complexity is given by the ratio between the lengths of the descriptions *D*_block_*(𝒯)* and *D*_list_*(𝒯)*.

**Figure S12.**
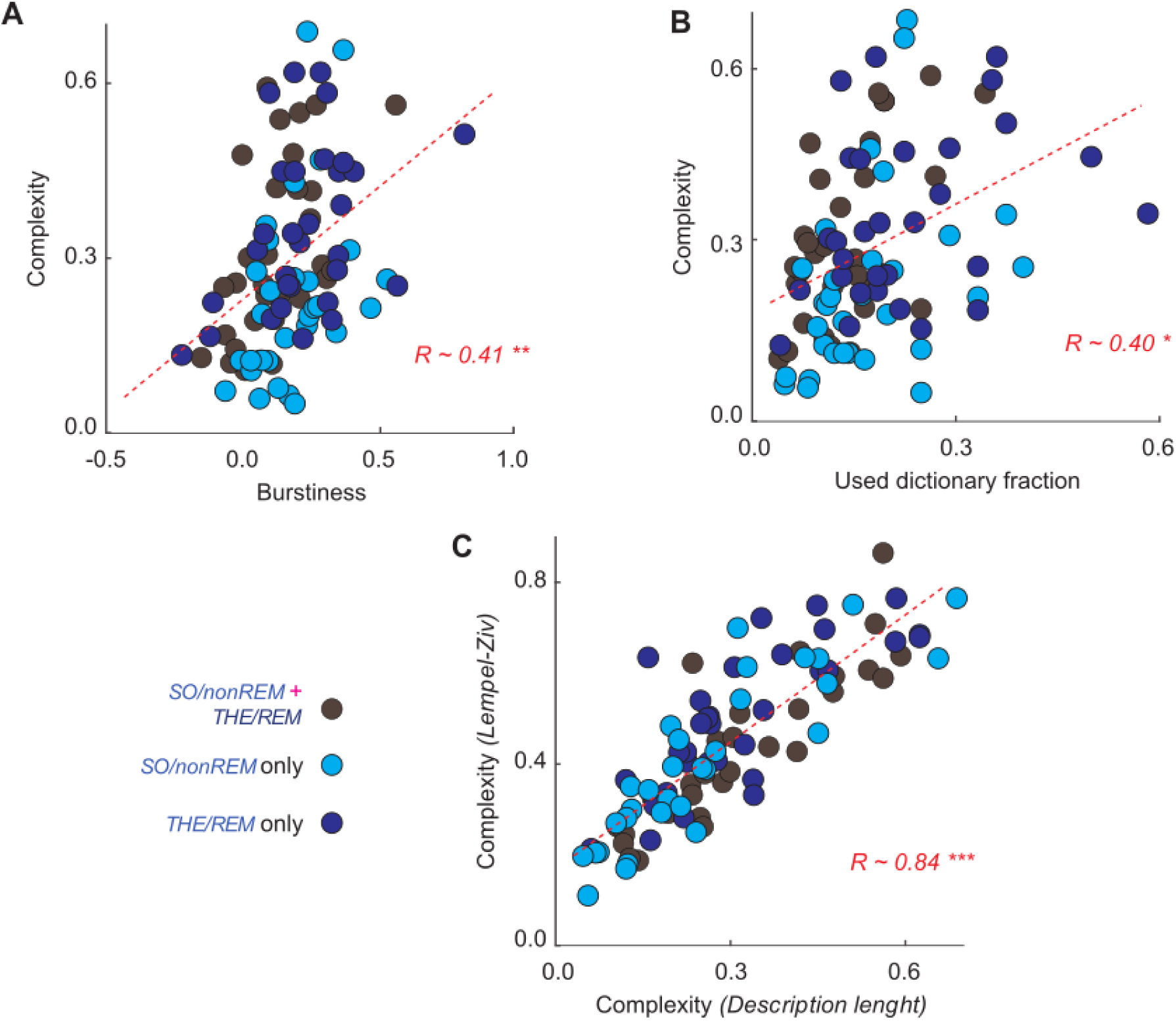
Burstiness and Used Dictionary Fraction explain complexity. Complexity linearly increased with burstiness **(A)** and Used Dictionary Fraction **(B)**. **(C)** Lempel-Ziv (LZ) complexity as a function of our measure of complexity. The two complexities were highly linearly correlated, and results of complexity analyses were thus qualitatively the same using either one of the two measures. We plot together results for complexity analyses restricted to SO/nonREM states (light blue dots), to THE/REM states (dark blue dots) or over all states combined (grey dots), as no significant differences were observed between the three groups with respect to the plotted linear trends.

**Figure S13.**
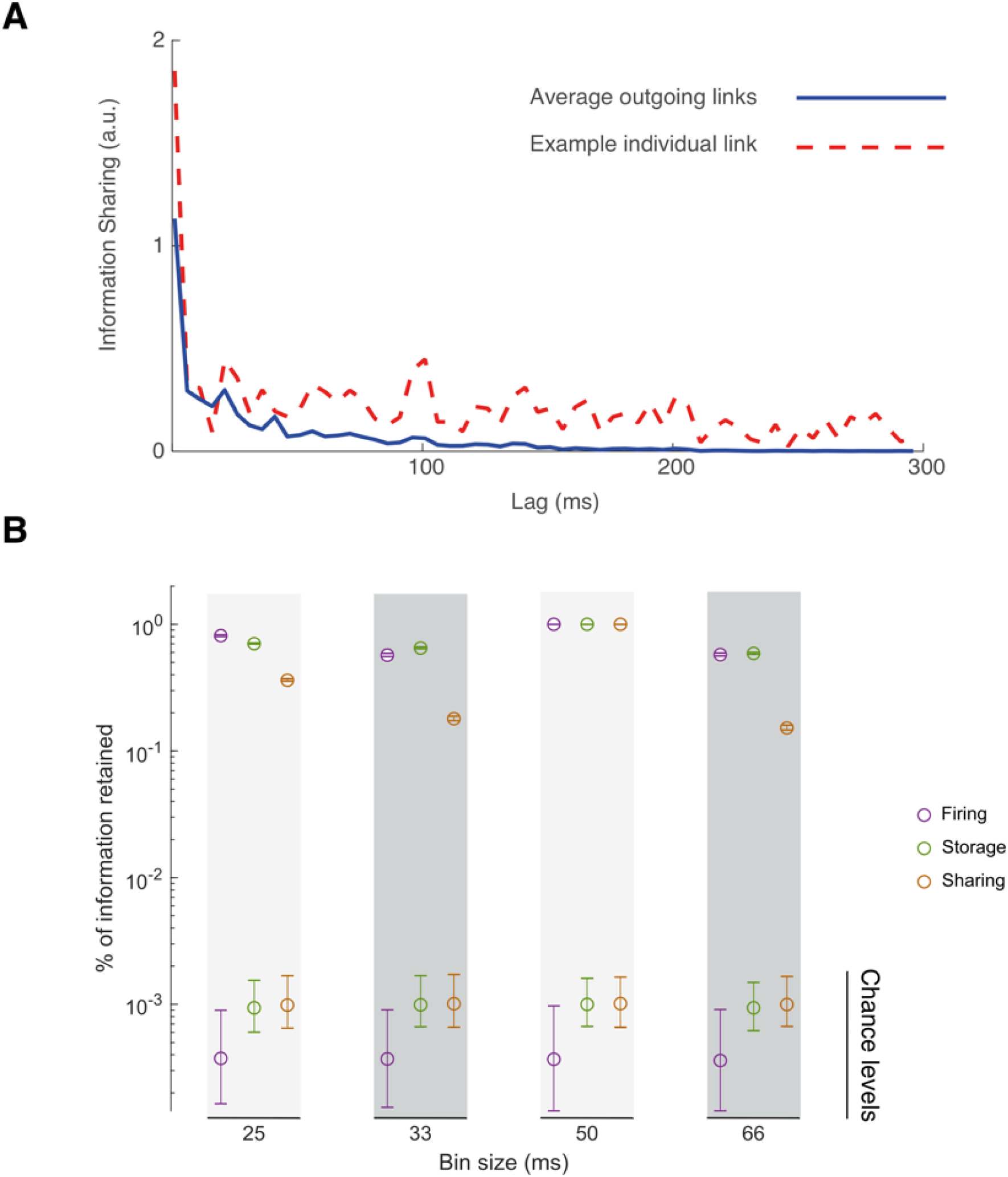
Additional robustness analyses. **(A)** Lagged mutual information terms between spike trains, the building block terms of both storage and sharing features, quickly decay as a function of the considered lag τ, justifying our choice to integrate MI only for latencies up to an average theta cycle period (∼125 – 250 ms depending on recordings). The red dashed line refers to a representative individual link, associated to a specific pair of units, and show some additional peak. However, these secondary peaks are much smaller than the main peak for very short latency and are not aligned for different pairs of units so that they are averaged out away when averaging over multiple outgoing links originating from a same unit (solid blue line). **(B)** Our procedure for substate extraction is robust against changes of the bin size. We considered four different choices of bin size different from the original choice of 50 ms, extracted substate sequences for each of these new bin choices and computed mutual information between the newly obtained and the original corresponding substate sequences. Shown here are the relative fraction of retained information for different bin sizes and substate types (evaluated on a representative recording, mPFC, natural sleep, the same as for figure S10). Across all features, the fraction of information retained about the substate sequences for the main reference bin size of 50 ms is two orders of magnitude above chance levels, with sharing being most effected. Vertical bars denote 99% confidence interval (bootstrap with replacement for original data, permutation-based for shuffled data, 1000 replicas in both cases).

## Supplementary tables

**Table S1.**
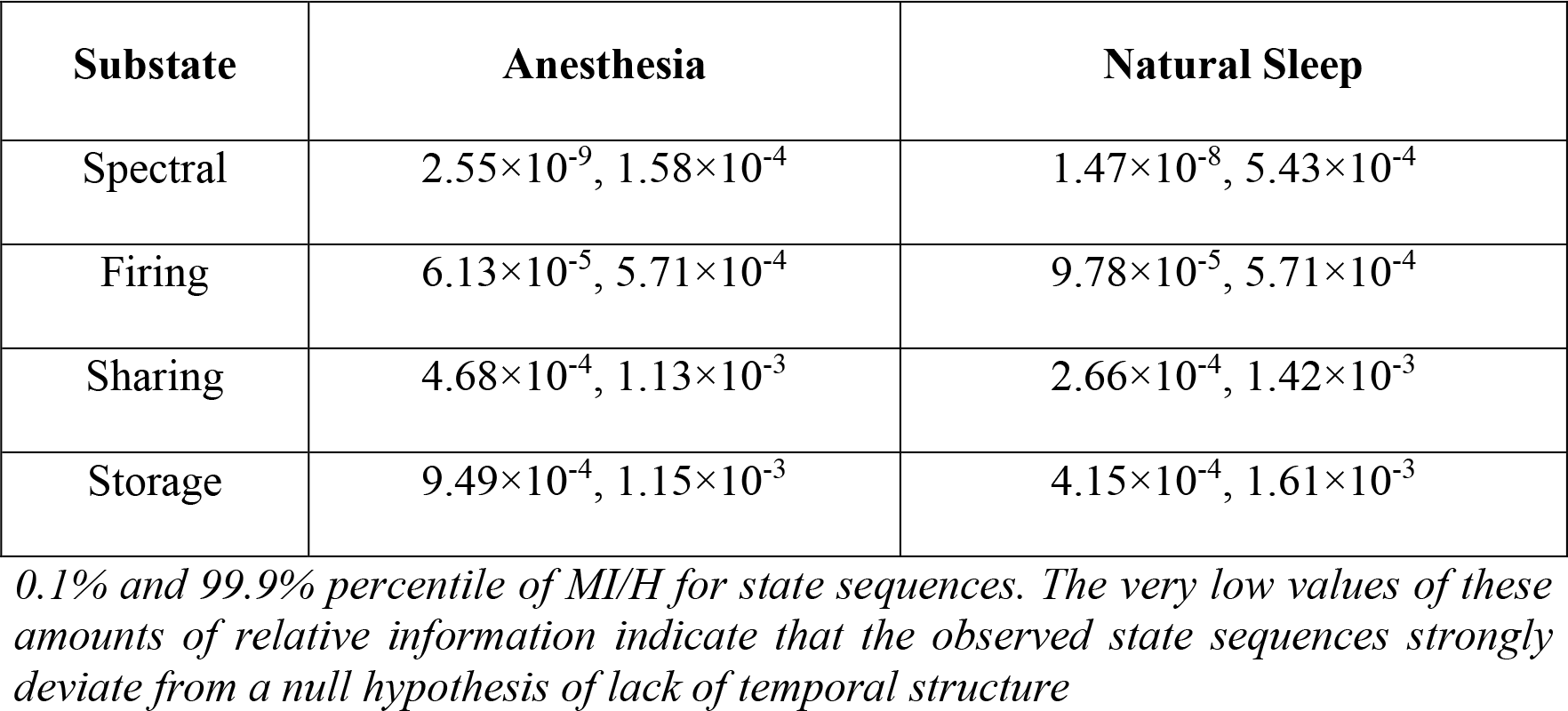
Percent of information about substate sequences in empirical recordings retained after shuffling

